# TASOR expression in naive embryonic stem cells safeguards their developmental potential

**DOI:** 10.1101/2024.02.25.581951

**Authors:** Carlos A. Pinzon-Arteaga, Ryan O’Hara, Alice Mazzagati, Emily Ballard, Yingying Hu, Alex Pan, Daniel A. Schmitz, Yulei Wei, Masahiro Sakurai, Peter Ly, Laura Banaszynski, Jun Wu

## Abstract

The seamless transition through stages of pluripotency relies on a delicate balance between transcription factor networks and epigenetic silencing mechanisms that ensure proper regulation of the developmental program, critical for normal development. Here, we uncover the pivotal role of the transgene activation suppressor (TASOR), a component of the human silencing hub (HUSH) complex, in sustaining cell viability during the transition from naive to primed pluripotency, despite its rapid downregulation during this transition. Loss of TASOR in naive cells triggers replication stress, disrupts H3K9me3 heterochromatin formation, and compromise the transcriptional and post-transcriptional silencing of LINE-1 (L1) transposable elements (TEs), with these effects become more pronounced in primed cells. Remarkably, the survival of *Tasor-*knockout cells during naive to primed transition can be restored through the inhibition of cysteine-aspartic acid protease (Caspase) or deletion of mitochondrial antiviral signaling protein (MAVS). This suggests that unscheduled L1 expression activates an innate immune response, leading to programmed cell death, specifically in cells exiting naïve pluripotency. Additionally, we propose that HUSH-promoted H3K9me3 in naïve PSCs sets the stage for ensuing DNA methylation in primed cells, establishing long-term silencing during differentiation. Our findings shed insights on the crucial impact of epigenetic programs established in early developmental stages on subsequent phases, underscoring their significance in the developmental process.

## Introduction

The naïve epiblast in the preimplantation blastocyst exhibits a transient state of global DNA hypomethylation due to epigenetic reprogramming following fertilization. Mouse and human pluripotent stem cells (PSCs) cultured in the presence of the MEK 1/2 inhibitor PD0325901 exhibit low DNA methylation akin to the preimplantation epiblast representing the naïve state of pluripotency^1–6^. Exiting the naive state can be triggered by removing MEK inhibition or via direct exposure to FGF2 and Activin A (FA), inducing mouse naïve PSCs to differentiate into transient formative epiblast-like cells (EpiLCs)^7^, which can be further stabilized in culture as primed epiblast stem cells (EpiSCs)^8,9^. EpiSCs resemble the post-implantation epiblast and are characterized by high levels of DNA methylation and an inactive X chromosome^10^.

The human genome is composed approximately 54% from repeat sequences that include more than 630 long interspaced nuclear elements (LINEs or L1s)^11^. In somatic cells, L1s are typically silenced through DNA 5mC CpG methylation^11^, yet in naïve PSCs, L1s are transcribed^12^ as a consequence of the hypomethylated genome^13^. The dysregulation of L1s has been linked to age-related disorders^11–1514^ and carcinogeneis^6,15^. Therefore, elucidating the mechanisms that safeguard the hypomethylated genome of naïve PSCs against the activation of repetitive elements could significantly enhance our understanding of the etiology of L1-associated disorders and may uncover new therapeutic interventions.

TASOR is a component of the human silencing hub (HUSH) complex, which is composed of two additional members, M-phase phosphoprotein 8 (MPP8) and periphilin 1(PPHLN1). The HUSH complex mediates gene silencing through H3K9me3 of repeats and intronless mobile elements^16^, particularly targeting the evolutionary young L1 endogenous TEs^17–20^. Despite previous studies, TASOR’s function in early embryonic development is not fully understood. Here, we dissect TASOR’s role in naïve pluripotency maintenance and exit through loss of function and epigenomic profiling studies.

### TASOR loss results in programed cell death upon the exit of naïve pluripotency

Many epigenetic features associated with the pre- and post-implantation epiblast, e.g. DNA methylation^21^, can be recapitulated by cultured PSCs (Figure S1A and S2A-S2B). To identify epigenetic regulators of epiblast development, we performed bioinformatic analyses using published datasets^22^ and compared chromatin interacting proteins during the transition from naïve mouse embryonic stem cells (ESCs) to formative epiblast-like cells (EpiLCs). We found that the HUSH component *Tasor* is highly expressed in naïve ESCs but rapidly downregulated at both the mRNA and protein levels upon differentiation (Figure 1A). We confirmed that TASOR protein is indeed lost in mouse (Figure 1B) and human (Figure S2C) primed PSCs. This observation is particularly interesting given previous reports showing that a loss-of-function mutation in *Tasor* leads to an *in vivo* gastrulation defect resulting in embryonic lethality^23,24^.

**Figure 1.**
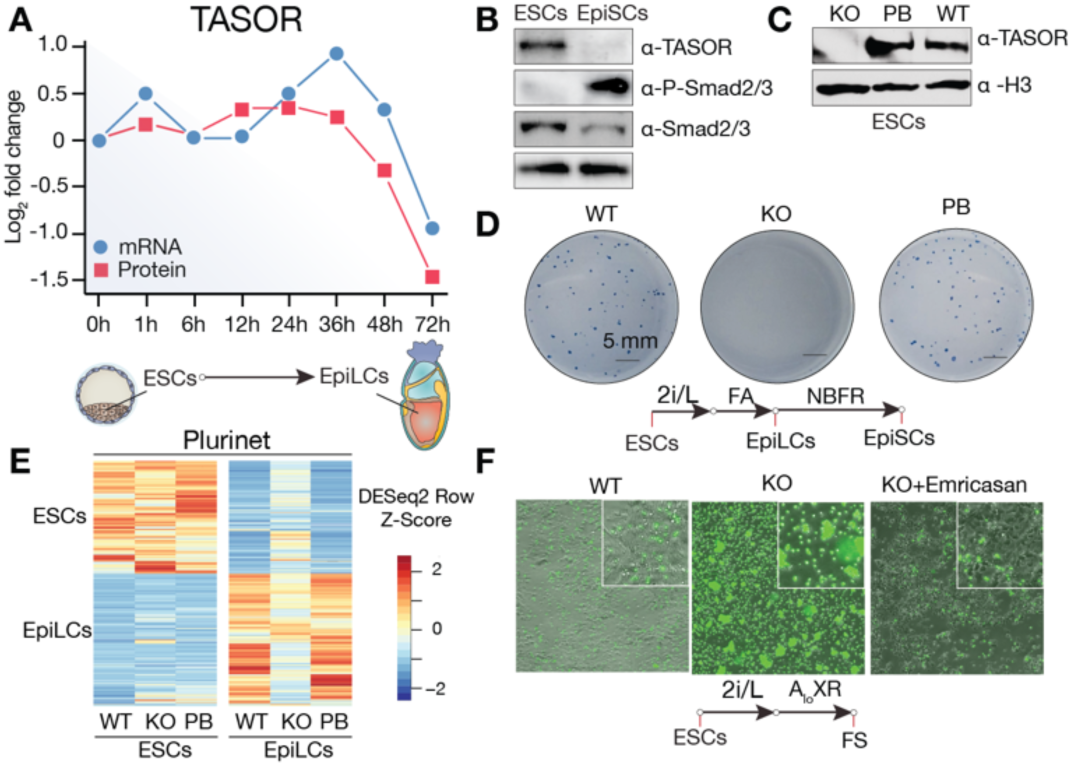
TASOR loss induces cell death during ESC differentiation. (**A**) Diagram of *Tasor* mRNA and protein levels in naïve ESCs to EpiLCs transition from the stem cell atlas dabase^22^. (**B**) Western blot of mouse embryonic stem cells (ESCs) cultured in 2i/L (Naïve), and epiblast stem cells (EpiSCs) cultured in NBFR (Primed). (**C**) Western blot for Wild type (WT), *Tasor* knockout (KO), putback (PB), and naïve ESCs cells cultured in 2i/L (**D**) Colony formation assay for naïve ESCs cells transitioned to EpiSCs. (**E**) Heatmap of different pluripotency markers between naïve ESCs to EpiLCs transition in from the plurinet dabase^105^. (**F**) SYTOX green cell death staining on cells transition to formative cells (FS) cultured for 72h in AloXR.

To investigate the role of TASOR in pluripotency, we generated *Tasor Knockout (*KO) naïve mouse PSCs (here after referred to as ESCs) using CRISPR-CAS9^25^ (Figures 1C and S1B-S1D). *Tasor-*KO ESCs maintained normal colony morphology and continued to express pluripotency markers when grown in the naïve (2i/LIF) condition (Figures S1E and S1F). We next tested how *Tasor* loss would affect the transition from naïve ESCs (2i/LIF) to formative PSCs conditions FAC (also known as FTW)^26^, and AloXR^27^. Surprisingly, although *Tasor* is drastically downregulated upon exit of the naïve state, we found that TASOR is required to establish formative PSCs, as only a few colonies are observed during the conversion in the absence of TASOR (Figures S1G and S1H). ESCs can transition to transient formative EpiLCs^7^ via exposure to FGF2 and Activin A (FA), and EpiLCs can be stabilized in culture as EpiSCs^26^ under FGF and WNT inhibition (NBFR). EpiSCs closely resemble the ectoderm of the late-gastrula-stage embryo^28,29^. Interestingly, we found that *Tasor* KO ESCs could transition to EpiLCs but failed to form colonies in the NBFR medium (Figure 1D). Notably, cellular differentiation phenotypes were rescued upon reintroduction of TASOR cDNA (TASOR putback, PB) in all conditions (Figures 1D, S1G and S1H). These findings suggest TASOR’s critical function in preserving cell viability after the transition from naïve pluripotency. Furthermore, they suggest that the gastrulation defect observed in *Tasor* mutant embryos *in vivo* could be partially due to the reduced survival of formative/primed epiblast cells.

We hypothesized that, in the absence of TASOR, the compromised *in vivo* differentiation potential might be due to an inability to properly establish differentiation-specific transcription programs. However, RNA sequencing (RNA-seq) revealed that, despite some transcriptional changes, *Tasor* loss was mostly compatible with differentiation to EpiLCs (Figure 1E), evidenced by the upregulation of *FGF5* and downregulation of *Nanog* expression (Figure S3A). Transcriptional changes were verified via qPCR for the upregulation of formative/primed marker genes *Otx2* and *Fgf5,* and the downregulation of *Prdm14* and *Tasor* (Figures. S3B-S3E). To assess the differentiation potential of *Tasor-*KO ESCs further, we performed teratoma analysis. Despite being smaller than those from WT controls, teratomas derived from *Tasor*-null ESCs contained tissues from all three germ layers (Figures S1I-S1K). Overall, these results indicate that despite compromised viability early in differentiation, ESCs lacking TASOR are still capable of differentiating into cells from all three primary germ layers.

Previously studies have demonstrated that p53 deficiency can partially mitigate the gastrulation defect observed in *Tasor* mutant embryos. This rescue effect is attributed to the reduction of apoptosis mediated by cysteine-dependent aspartate-specific proteases (CASPASE)^24,30^. To determine whether the diminished colony-forming capacity of *Tasor-*KO ESCs upon transitioning to formative and primed states was due to cell death, we used SYTOX green staining to quantify dead and dying cells following formative (AloXR) transition. In contrast to WT cells, *Tasor-*KO resulted in extensive cell death during the transition (Figure 1F). Cell death can occur through several mechanisms, including apoptosis, pyroptosis, and necroptosis^30^, or through a combination of these pathways known as PANoptosis^31^. With the main shared mechanism for some of these pathways being by the cleavage of cysteine-dependent aspartate-specific proteases (CASPASE). To determine the mode of cell death in *Tasor-*KO cells, we applied the pan-CASPASE inhibitor Emricasan and found that 5 µM Emricasan significantly restored the colony-forming capability and greatly enhanced cell survival following formative transition (Figure 1F). This evidence, along with findings from prior studies, suggests apoptosis as the primary mechanism of cell death, though the potential contribution of pyroptosis or PANoptosis has not been conclusively excluded. These findings collectively indicate that TASOR is crucial for the transition from naïve to formative state, with the loss of *Tasor* in ESCs leading to marked increase in programmed cell death among formative cells.

### TASOR loss leads to increased DNA damage and cell cycle arrest

Previous studies have demonstrated that disrupting the HUSH complex leads to DNA damage and cell cycle arrest^32,33^. Consistently, the loss of TASOR resulted in a longer doubling time, which was reversed upon TASOR reintroduction (Figure S4A). Cell cycle analysis via EdU incorporation revealed that the increased doubling time is due to both an elongated G2/M phase and a shortened S phase (Figures 2A and S4B). Additionally, there was a decrease in the number of cells positive for the mitotic marker phospho (Ser 10) H3 (Figures 2B, S4C and S4D), indicating that the accumulation cells in G2/M phase is likely caused by a G2 arrest. We investigated whether DNA damage might contribute to the observed G2 arrest. Our findings revealed that the absence of *Tasor* is associated with elevated levels of phospho(S139) gamma H2AX (γ-H2AX) (Figure 2C), along with increased phospho p53, and phospho DNA-PKcs (s2056) (Figure S4E). Additionally, RNA-seq indicated an increase in the expression of retinoblastoma (*Rb1*) in *Tasor* KO ESCs (Figure S4F). Notably, both *Rb1* expression levels was further upregulated upon transition to EpiLCs with the addition of the cyclin-dependent kinase inhibitor 1A (*Cdkn1a*, also known as p*21*) (Figure S4G), suggesting that enhanced activation of p53-p21-Rb pathway^34^ might be driving the G2 cell cycle arrest.

**Figure 2.**
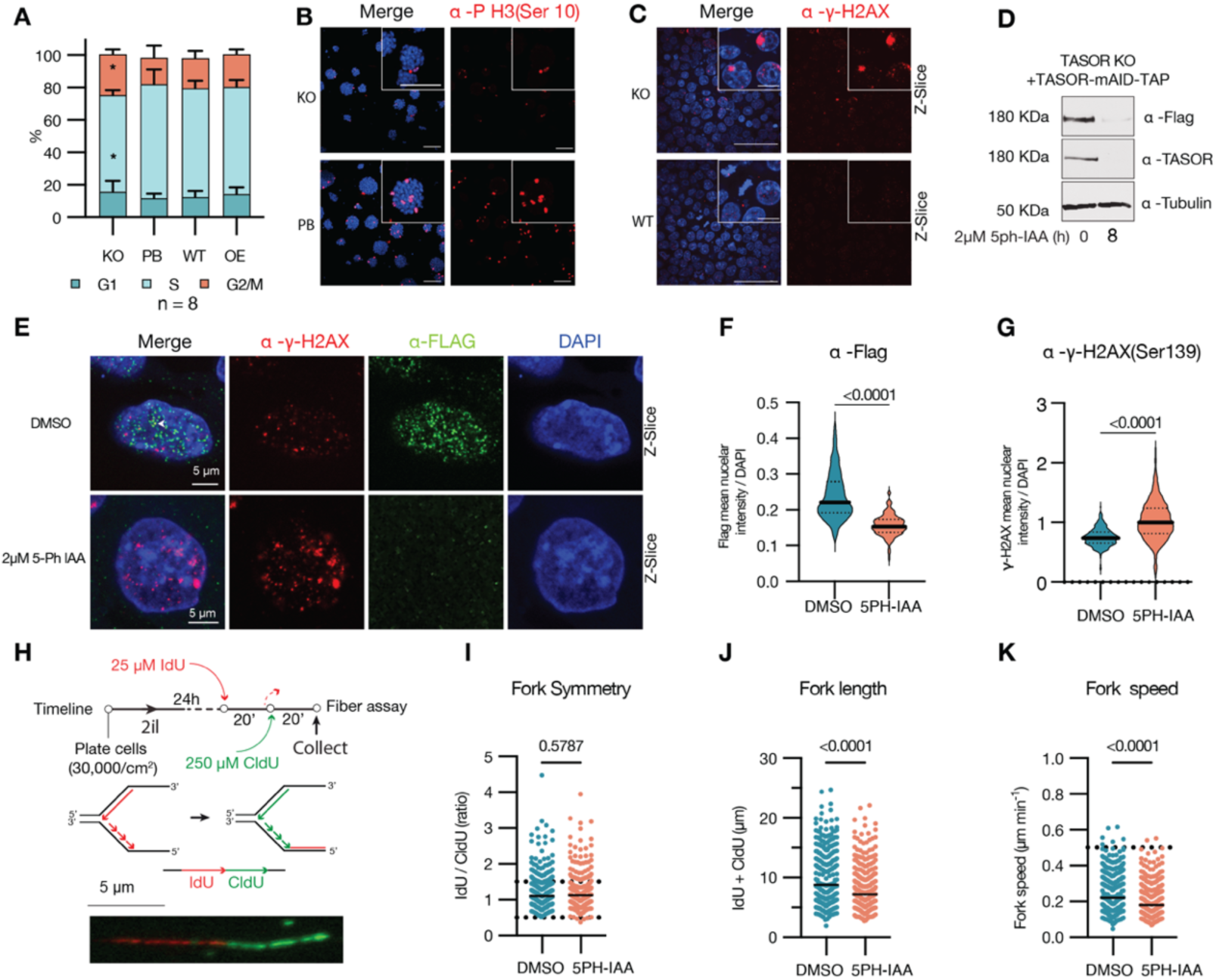
TASOR loss induces cell cycle arrest, DNA damage and DNA replication stress. (**A**) Flow cytometry cell cycle analysis via Edu incorporation and DNA staining of *Tasor* KO, PB, WT, and overexpression (OE) naïve ESCs cultured in 2i/L. (**B**) Immunostaining for the mitosis marker phospho H3 (Serine 10) for *Tasor* KO and PB naïve ESCs in 2i/L. (**C**) Immunofluorescence staining of WT and *Tasor* KO cells for phosphoserine 139 of histone H2AX (γH2AX). (**D**) Western blot for TASOR 8 hours after addition of 2µM 5ph-IAA. (**E**) Z-slice confocal immunofluorescence image for γH2AX in control DSMO or 5-ph-IAA treated cells. (**F**) Mean segmented nuclear intensity normalized to DAPI of FLAG Alexa fluor 488. (**G**) Mean segmented nuclear intensity normalized to DAPI of γH2AX Alexa fluor 555. (**H**) Diagram of DNA fiber assay with representative image of chromatin fiber. (**I**) Replication fork symmetry quantification of chromatin fibers in DMSO or 5-ph-IAA treated cells (n=2). (**J**) Replication fork length quantification of chromatin fibers in DMSO or 5-ph-IAA treated cells (n=2). (**K**) Replication fork speed quantification of chromatin fibers in DMSO or 5-ph-IAA treated cells (n=2).

To elucidate the immediate impact of TASOR loss on DNA damage, we engineered an ESC line containing Oryza sativa TIR1 (OsTIR1) F74G and a *Tasor* re-expression vector with a C-terminal mini auxin-inducible degron (mAID)^35,36^. Treatment with 2 µM 5-phenyl-indole-3-acetic acid (5-Ph-IAA) rapidly degraded TASOR (Figures 2D and S4H), resulting in a marked decrease in the nuclear signal of TASOR-flag (Figures 2E and 2), H3K9me3 (Figure S4J), phospho-H3 (Figure S4L), and MPP8 (Figure S4M), while leaving H3K27me3 levels unchanged (Figure S4N). These observations suggest that the acute loss of *Tasor* mirrors the effects of its chronic absence. Using this system, we confirmed a significant increase in γ-H2AX foci 24 hours post-TASOR depletion (Figures 2E and 2G). Further analysis through co-staining for FLAG-tagged TASOR, H3K9me3 and γ-H2AX, revealed TASOR appeared as nuclear punctuate and co-localized with some H3K9me3 and γ-H2AX foci (Figures 2E, S4I-S4J). Given that H3K9me3 accumulates at stalled replication forks and DNA double-strand breaks^37,38^, and its loss can compromise replication fork stability^39^ and double stranded break repair^38^, we determined whether the increased γ-H2AX signaled replication fork instability. By labeling replication tracts with 5-chloro-2′-deoxyuridine (CldU) and 5-iodo-2′-deoxyuridine (IdU) and analyzing DNA fibers (Figure 2H), we found that TASOR depletion reduced both replication fork length and speed, without affecting fork symmetry (Figures 2I-2K), pointing to replication fork stress in the absence of TASOR.

### Interplay between TASOR and DNA methylation in naïve ESCs

To study the link between DNA methylation and TASOR’s role in H3K9me3 regulation, we adjusted the global DNA methylation levels in ESCs. MEK inhibition leads to a dose-dependent decrease in DNA 5mC methylation in ESCs, a process attributed to the loss of UHRF1 and the reduced expression of the *de novo* methyltransferases Dnmt3a, Dnmt3b and Dnmt3l^40,41^. This reduction is facilitated by the transcription factor PRDM14^42^. Consequently, lowering the MEK inhibitor PD0325901concentration from 1 µM to 0.2 µM in titrated 2i/L (t2i/L) increases global DNA methylation levels without altering the pluripotency state^43^. Conversely, the addition of L-ascorbic acid (a form of Vitamin C) promotes DNA hypomethylation, as vitamin C acts as a cofactor for Fe(II)/2-ketoglutarate-dependent (Fe/αKG) dioxygenases by reducing Fe^3+^ to Fe^2+^ ^44,45^. Fe/αKG dioxygenases include various epigenetic regulators like Jumonji C domain-containing histone demethylases (JHDMs, e.g., KDM4A/C), DNA and RNA demethylases (ALKBH family), and the TET family of DNA hydroxylases^44,46,47^. Thus, adding L-ascorbic acid can lead to reductions in DNA 5mC^48–50^, H3K9me3^51^, and m6A RNA levels^52,53^(Figures S1A, S5A and S5B).

Compared to controls (WT, PB and OE ESCs), we found *Tasor-*KO ESCs were more vulnerable to global hypomethylation induced by vitamin C, resulting in a more pronounced increase in doubling time and cell cycle arrest (Figures S5C and S5D). The addition of 100 µg/ml of vitamin C^4,50^ further exacerbated the derepression of L1 ORF1 protein levels (Figure S5B) and led to an increase in segregation errors and chromosomal abnormalities (Figures S5F-S5L). Interestingly, when cultured in t2i/L, both WT and *Tasor-* KO ESCs showed an increase in global H3K9me3 levels (Figure S5B) and a decrease in doubling time compared to those cultured in standard 2i/L condition (Figures S5C and S5E). These findings suggest partial rescue of *Tasor-*KO ESCs phenotypes when cultured with reduced MEK inhibition^54^. A similar compensatory response was noted in *Mpp8-*KO ESCs grown in the serum/LIF condition^54^, which might be facilitated by a polycomb-mediated epigenetic switch involving H3K27me3^50^.

To determine the sequence of events involving DNA methylation and TASOR-mediated H3K9me3, we compared *Tasor-*KO ESCs with triple *Dnmt* knockout (*Dnmt*-3xKO) ESCs, which lack *Dnmt1*, *Dnmt3a*, and *Dnmt3b*. Immunostaining revealed that *Dnmt*-3xKO ESCs exhibited a loss of DNA methylation marks 5mC and 5hmC (Figures S6A and S6B), yet retained histone modifications H3K9me3 and H3K27me3 (Figures S6C and S6D). Notably, *Dnmt*-3xKO ESCs mostly did not phenocopy *Tasor* loss, including the prolonged doubling time and decreased number of cells positive for the phospho (Ser 10) H3 (Figures S6E and S6F). Additionally, upon culturing in AloXR for 48h, we did not observe substantial cell death (Figure 4C), consistent with a previous report^55^. However, *Dnmt*-3xKO ESCs exhibited a similar increase in L1 transcripts as observed in *Tasor-*KO ESCs (Figure 4B). These results suggest that in naïve ESCs, both TASOR-mediated H3K9me3 and DNA methylation are critical for L1 repression. This is in contrast to what is observed in somatic cells where DNA methylation plays a more essential role, which is evidenced by the observation that upon TASOR loss, L1 elements remain repressed through strong promoter methylation^56,57^.

### TASOR regulates steady-state LINE1 RNA transcripts

HUSH complex is well known for its role in L1 retrotransposon silencing^58^. To investigate whether loss of retrotransposon silencing contributes to cell death in *Tasor-*KO ESCs during the transition to the formative state, we first mined our RNA-seq data. Our analysis revealed that TASOR loss led to a significant increase in the steady-state levels of L1 RNA in ESCs, particularly within the evolutionarily young L1MdTf family of LINEs (Figure 3A). Upon transition to EpiLCs, the absence of TASOR resulted in an even more pronounced surge in LINE transcript abundance (Figure 3B), which was further confirmed by immunostaining (Figure S1O). During the formative transition, the L1MdTf subfamily was the most affected, with notable dysregulation also observed in the L1MdA and L1MdG subfamilies (Figures 3B and 3C). Beyond LINEs, modest yet significant increases in transcript abundance were observed for two endogenous retrovirus (ERV2) subfamilies, MMETn and ETnERV, as well as for several satellite repeat subfamilies, including general satellites (GSAT_MM), centromeric satellite (CENSAT_MM), and minor satellite repeats (SYNREP_MM). These satellite families exhibited substantial fold changes to WT, albeit at relatively low transcript levels (Figures S7A and S7F-S7I). Given the most striking changes in transcript abundance observed in *Tasor*-KO EpiLCs for LINEs, our further analyses focused on these elements.

**Figure 3.**
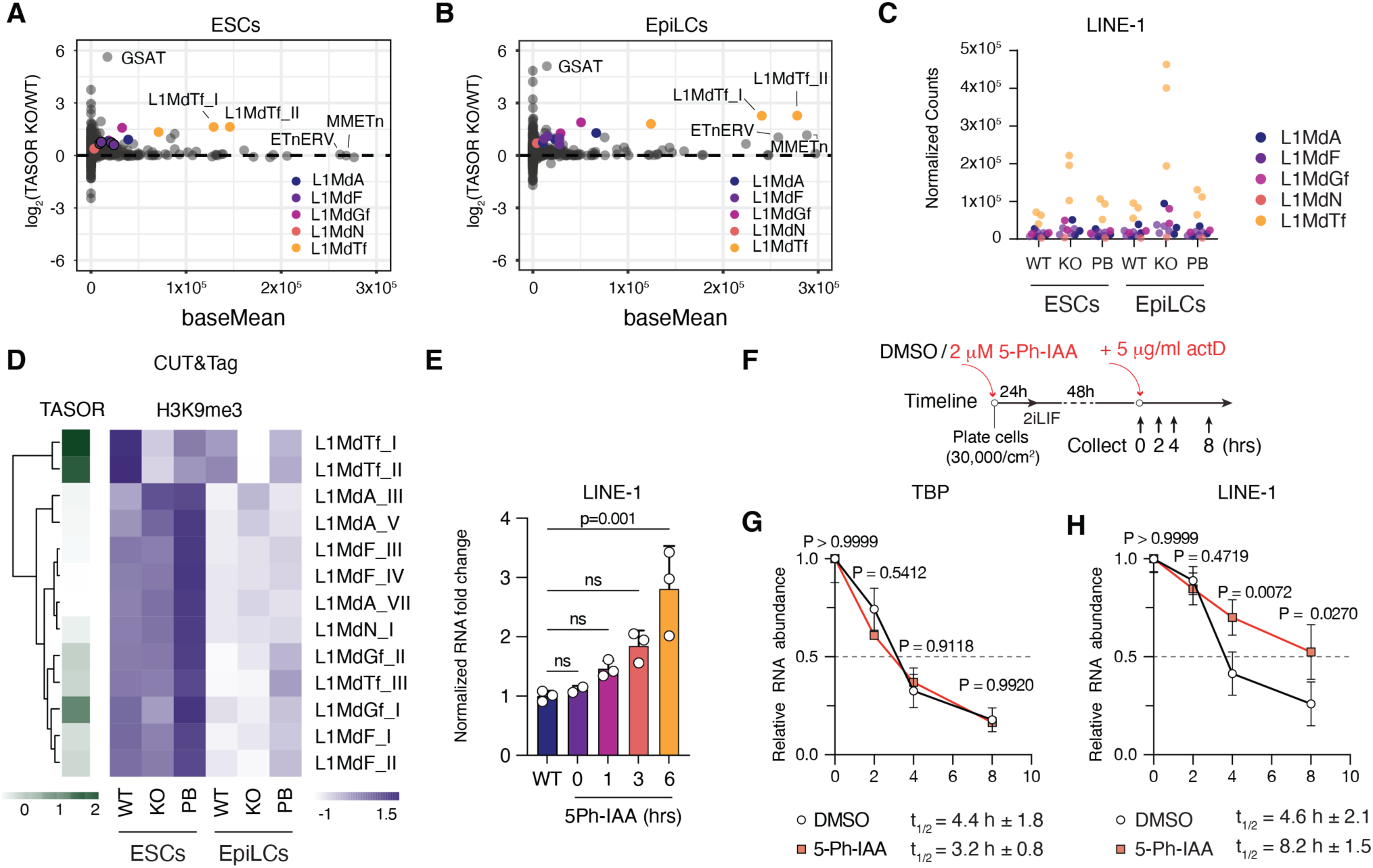
L1 RNA abundance and half-life is increased upon TASOR loss. (**A**) MA plots for repeats showing log2 fold change of Tasor *KO* (KO) over *Wildtype* (WT) in naïve ESCs. (**B**) MA plots for repeats showing log2 fold change of Tasor *KO* (KO) over *Wildtype* (WT) in EpiLCs. (**C**) Normalized average counts for LINE-1 (L1) sub families in naïve ESCs and EpiLCs for WT, KO and Tasor Putback (PB). (**D**) Heatmap for CUT&Tag of TASOR-3xFlag and H3K9me3 at L1 family members. (**E**) Timecouse normalized RNA fold change via qPCR for L1 after Auxin treatment. (**H**) Experimental diagram for measuring RNA half live after Actinomycin D (ActD) treatment. **(I**) Relative mRNA abundance after ActD treatment for TBP measured by RT-qPCR. (**J**) Relative mRNA abundance after ActD treatment for L1 measured by RT-qPCR.

The HUSH complex has previously been associated with the silencing of LINE elements through the deposition of H3K9me3^17,18,59^. To investigate the impact of TASOR loss on H3K9me3 levels at LINE elements, we performed Cleavage Under Targets and Tagmentation (CUT&Tag)^60^ for Flag (TASOR) and H3K9me3. Our analysis revealed that TASOR predominantly binds to the 5’ regulatory region of L1MdTf elements, with lesser binding observed across various other L1 subfamilies (Figures 3D and S7B). In comparison, H3K9me3 was highly enriched across a broad spectrum of repetitive elements, including LINEs, ERVs, telomeric and satellite repeats (Figures 7F and 7H), with H3K9me3 peaks at LINE elements frequently located near the 5’ end (Figure S7C). Following *Tasor* KO, a notable reduction in H3K9me3 was observed in both ESCs and EpiLCs, especially at L1MdTf subfamilies (Figures 3D and S7C). This decrease in H3K9me3 at L1MdTf elements was reversed upon reintroduction of Tasor cDNA (Figure S7C). Furthermore, immunostaining, flow cytometry and immunoblot analyses also revealed a partial reduction in H3K9me3 levels in both mouse and human TASOR^-/-^ ESCs (Figures S1L-S1N, and S2I-S2J). These results demonstrate TASOR’s critical role in establishing and/or maintaining H3K9me3 at the 5’ end of specific LINE subfamilies, and highlight that the loss of TASOR and subsequent reduction of H3K9me3 are linked to increased LINE RNA abundance.

Since H3K9me3 is closely linked to gene silencing and predominantly found in heterochromatin, we explored whether the loss of H3K9me3 at L1MdTf elements would result in increased chromatin accessibility. To this end, we performed an assay for transposase-accessible chromatin followed by sequencing (ATAC-seq)^61^. Contrary to the significant reduction of H3K9me3 at L1MdTf elements following *Tasor* KO, the increase in ATAC signal in *Tasor*-KO ESCs and EpiLCs was only modest (Figure S7E). Interestingly, the transition of WT cells from ESCs to EpiLCs resulted in both a higher ATAC signal and a decrease in H3K9me3 at L1MdTf, without altering L1 RNA levels (Figures 3D and S7C-S7E). This observation suggests that during this transition, alternative silencing mechanisms, such as DNA methylation might come into play.

Motif analysis of ATAC peaks revealed that upon transition to EpiLCs, WT ESCs showed reduced accessibility at motifs associated with pluripotency transcription factors (SOX2, POU5F1) and increased accessibility at motifs associated with DNA methylation and imprinting (ZFP57), and higher-order chromatin organization (CTCF) (Figure S8A). In contrast, *Tasor*-KO ESCs exhibited reduced accessibility at ZFP57 binding sites relative to WT cells (Figure S8B). Furthermore, during the transition, *Tasor*-KO cells failed to decommission transcription factors from the KLF and POU families, including POU5F1 (also known as OCT4) (Figure S8C). These results suggest that although *Tasor*-KO ESCs could differentiate to EpiLCs, some transcriptional differences can be due to the inability to properly decommission ESC-specific transcription networks (Figure 1E).

Interestingly, we noted the transcriptional dysregulation of various imprinted genes (Figure S8D), including notable changes in *Igf2* (Figures S8E-F), and observed that certain genes containing internal L1s were derepressed in a manner dependent on the orientation of the L1 sequence (Figure S8G). Specifically, genes such as *Mrc1 and Fsd1l* exhibited derepression only in exons downstream of the L1 elements (Figures S8H and S8I). Additionally, during the transition of *Tasor*-KO ESCs to EpiLCs, there was an upregulation in the expression of gene exons adjacent to L1 sequences (Figure S8J). This observation can potentially be explained by recent reports of L1s acting as “gene traps” during splicing events, leading to the creation of chimeric transcripts^62,63^, or due to the disruption of H3K9me3 “spreading” mediated by the HUSH complex and MORC2 at these loci^64,65^.

Beyond its involvement in H3K9me3 deposition, the HUSH complex has also been implicated in the targeted degradation of L1 RNA via interactions with the nucellar exosome targeting (NEXT) and CCR4-NOT complexes^16,66–68^. To study the effect of TASOR loss on L1 RNA stability, we utilized the auxin-inducible TASOR-mAID line. Treatment with auxin led to a rapid increase in L1 transcripts, coinciding with the depletion of TASOR protein (Figures 3E, and 2D-2F). Using actinomycin D to halt further transcription, we tracked the persistence of selected RNAs over 8 hours in the presence or absence of TASOR (Figure 3F). In control cells, the half-life of transcripts not typically targeted by be HUSH complex, such as TBP, remained unchanged (Figure 3G). Strikingly, auxin-induced TASOR depletion significantly increased the half-life of L1 RNA (t_1/2_ = 8.2h ±1.5) compared to the DMSO-treated control (t_1/2_ = 4.6h ± 2.1) (Figure 3H), indicating that the observed increases in L1 RNA following TASOR loss may be partly attributed to decreased degradation of L1 transcripts.

### Cell death upon *Tasor*-KO ESC-to-EpiLC transition is mediated by innate immunity

The HUSH complex plays a pivotal role in linking retrotransposon silencing with innate immunity, primarily through its control of L1 expression and the subsequent MAVS-dependent sensing of L1 RNA*^18^*^69^. In the context of cancer, disruptions to the HUSH complex leading to L1 dysregulation have been shown to trigger innate immune pathways, resulting in the death of cancer cells^19,69^. We speculated whether the cell death observed during the transition from *Tasor*-KO ESCs to EpiLCs could be due to an innate immune response. To test this, we first generated *Mavs* and *Tasor* double KO ESCs (Figure 4A), which exhibited somewhat reduced IRF3 dimer formation compared to *Tasor* KO (Figure S9A). Surprisingly, viability assessments after 48 hours in the AloXR condition revealed that the loss of MAVS nearly completely rescued the cell death seen in *Tasor*-KO EpiLCs (Figure 4C). Flow cytometry analysis confirmed that *Mavs* and *Tasor* double KO decreased the percentage of dead cells to levels simiar to WT (Figures 4C and 4D). However, when cells were further transitioned from formative EpiLCs to primed EpiSCs, we observed diminished colony formation and alkaline phosphatase (AP) staining when compared to WT or PB cells, though the outcomes were still an improvement over *Tasor* KO (Figure S9B). Surprisingly, *Mavs* and *Tasor* double KO ESCs demonstrated a significantly reduced capacity to form teratomas compared to *Tasor* KO alone (Figures S9C-S9D), suggesting that while *Mavs* KO mitigates cell death immediately following exit from naive pluripotency, its simultaneous loss with TASOR introduces a synthetic lethality in later developmental stages. Unexpectedly, two separate *Mavs* and *Tasor* double KO clonal lines both showed markedly increased levels of L1 RNA when compared to *Tasor* KO alone (Figure 4B). Considering L1 RNA’s role in activating MAVS-mediated innate immune pathway^19,69^, this increase might stem from a subpopulation of cells that highly express L1, which would typically undergo apoptose in *Tasor* KO but survive in *Mavs* and *Tasor* double KO lines. Taken together, these results show that while cell death at the formative EpiLC stage is MAVS-dependent, other mechanisms likely contribute to later developmental stages.

**Figure 4.**
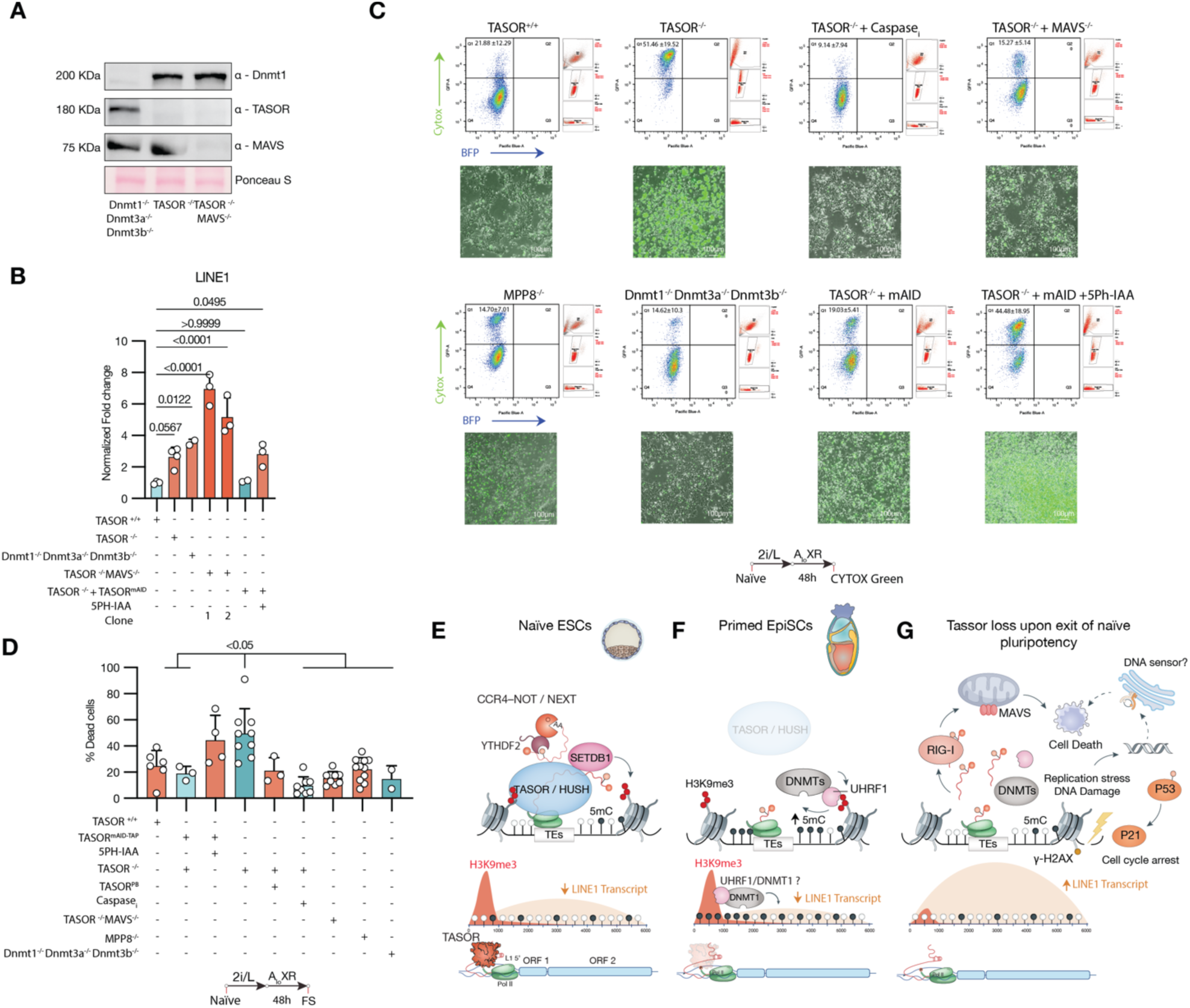
TASOR loss induced cell death is partially mediated by a MAVS innate immune response. (**A**) Western blot for DNMT1, TASOR and MAVS for Dnmt triple knockout (KO, *Dnmt1, Dnmt3a, Dnmt3*), *Tasor* KO, TASOR and MAVS double KO cells. (**B**) RT-qPCR for L1abundace in Wild type, *Tasor* KO, *Dnmt*3xKO, *Tasor* and *MAVS* double KO cells, and TASOR mAID with and without 2µM 5ph-IAA treatment. (**C**) Representative epifluorescence images for cell death via SYTOX green staining (bottom) and flow cytometry analysis (top) after 48 hours of AloXR formative cell conversion. (**D**) Flow cytometry quantification of the percent of dead cells measured via SYTOX green dead cell staining. (**E**) Proposed model diagram: in naïve ESCs, L1s and repeats are transcribed by Pol II, the nascent RNA transcript gets recognized by HUSH, which then recruits SETDB1 for depositing H3K9me3 with the help of the ATPase chromatin remodeler (MORC2) (not shown), the RNA transcript is marked by m6a, recognized by YTHDC1 and the RNA is targeted for degradation by the NEXT and the CCR4-CNOT complex. (**F**) In primed cells, TASOR is downregulated, and the H3K9me3 marked sites get targeted for 5mC CpG DNA methylation and long-term silencing. The UHRF1 Dnmt1 read-and-write model for the H3K9me3 and 5mC cross talk is depicted. (**G**) In *Tasor* KO ESCs upon exit of naïve pluripotency L1 transcripts are recognized possible by a RIG-I sensor and initiates a MAVS mediated “innate immune checkpoint”. The observed DNA damage and P53-P21-Rb mediated cell cycle arrest is depicted, as well as a possible DNA stimulated innate immune response.

## Discussion

Our results demonstrate that TASOR/HUSH plays an important role in establishing H3K9me3-mediated heterochromatic silencing at L1 sequences and repeats in naïve PSCs. One possible mechanism to achieve this is by recognizing the nascent RNA transcript and targeting them for degradation similar to the yeast homolog of HUSH^20^, the RNA-induced transcriptional silencing (RITS) complex^70–72^. By ensuring no productive transcripts are produced may explain the observed phenomenon in naïve PSCs, where transcription of L1s contributes to chromatin accessibility^12^ and enhancer formation^73^, yet retrotransposition of L1s remains low.

H3K9me3 is essential for initiating DNA CpG methylation and maintaining low levels of histone acetylation, both key features of heterochromatin^74–76^. Our results suggests that the deposition of H3K9me3 in the preimplantation epiblast is crucial for setting the stage for DNA methylation post-implantation^74,75^, with these methylation patterns being preserved throughout development to ensure the long-term repression of repetitive elements and stabilization of cell fates^77^. It is likely that multiple mechanisms coordinate this H3K9me3–5mC crosstalk. In a simplified read-and-write model, UHRF1^78^ reads H3K9me3 signals to guide DNMT1, thereby facilitating DNA methylation at H3K9me3 sites. DNMT1 can also directly interact with H3K9me3 through its tandem tudor domain (TTD)^78,79^, while the MPP8 chromodomain links the histone methyltransferases GLP/G9a with methylated DNMT3A, facilitating their interaction^80^. Additionally, protein-protein interactions between G9a-GLP and Dnmt3a further support this regulatory network^74^.

Our findings demonstrate that TASOR loss affects L1 mRNA stability and show that acute TASOR depletion triggers replication stress. Hush is known to interact with the co-transcriptional termination machinery ^68^, the RNA deadenylase CCR4-CNOTcomplex scaffold protein CNOT1^66^, and components of the nucellar exosome targeting (NEXT) complex^81^. Increased L1 mRNA stability has been observed following the KO of the m6A demethylase FTO^82^, and HUSH is known to interact with the m6A reader YTHDF2^66,67^. While some researchers advocate for Mettl3^83^ as the m6A writer involved in this process, others argue that Mettl3 does not specifically target L1s for degradation^84^.

A recent paper shows that in serum/LIF-cultured ESCs, the HUSH complex interacts with the leading strand DNA polymerase ε (POLE) complex, specifically the POLE1/2 subunits. This interaction promotes the asymmetric transfer of H3K9me3 to the leading strand of the replication fork, and this H3K9me3 asymmetry silences “head on” orientation L1 expression in the S phase of the cell cycle ^33^. The loss of POLE1/2 or TASOR can lead to increased DNA damage, as indicated by γH2AX^33^, suggesting this interaction is crucial for preventing replication stress and DNA damage. Additionally, genome wide profiling of R-loop in ESCs^85^ shows R-loops can accumulate at L1 5’ promoters (Figure S8G). This R-loop accumulation at L1 promoters during DNA replication may be a causal determinant of DNA damage, as it could cause collisions with the replication forks ^86–88^. Interestingly, the RNA–DNA helicase *DHX9* plays a protective role by unwinding of R-loops and G-quadruplexes^89^, and its loss results in defective H3K9me3 heterochromatin inheritance^90^.

The observed increase in chromosome segregation errors, micronuclei, and abnormal karyotype (Figure S5) indicates that TASOR loss may lead to genomic instability through mechanisms beyond replication stress. The presence of TASOR at major satellite and centromeric repeats, together with the reduction of H3K9me3 following TASOR depletion and the concurrent gain of major satellite repeat RNA, as evidenced by RNA-seq and qPCR analyses, points to a potential effect on centromere integrity. This notion is supported by studies involving epigenetic remodeling through a TALE-demethylase targeted to centromeric repeats^91^, which have shown a reduction in H3Kme3 levels, impaired HP1 recruitment, and subsequent impacts on chromosome segregation during mitosis.

ESCs are immunologically different from somatic cells^92^. Unlike differentiated cells, ESCs exhibit a much weaker response to cytoplasmic double-stranded RNA (dsRNA) and produce minimal amounts of IFN-β^93^. This reduced innate immune response might serve as a protective mechanism, allowing ESCs to avoid immune-related cytotoxicity^92^. Our results suggest that the inability to silence ERVs and repetitive elements activates an “innate immune checkpoint”, leading to the elimination of cells that cannot suppress these elements during naïve-to-primed transition. This hypothesis is supported by findings that depletion of *Tasor*^23^*, YTHDC1*^94^, SETDB1^95–97^, G9a/GLP^98,99^, SUV39H1/2^99–101^*, Mettl3*^94^, or *Dicer*^102^ results in L1 derepression and embryonic lethality. However, mice lacking FTO^103^ or MPP8^32^ are viable, likely due to compensatory mechanisms, as evidenced by increased TASOR binding in the absence of MPP8^18,33^.

Our study reveals that TASOR depletion impedes the proper transition of ESCs out of naïve pluripotency in a MAVS-dependent manner, resulting in CASPASE-dependent apoptosis. This cell death can be mitigated by Emricasan. Furthermore, TASOR loss triggers a P53-P21-Rb mediated DNA damage response and G2 cell cycle arrest, a phenomenon similarly observed in cells^87,88^ and cancer models where L1 retrotransposon significantly impacts cancer growth and progression^32,87,104^.

In conclusion, our findings underscore TASOR’s critical role in the maintenance and exit of naïve pluripotent state and reveal the intricate relationship between epigenetic regulation and innate immunity during embryonic development.

---

**Supplementary Figure 1.**
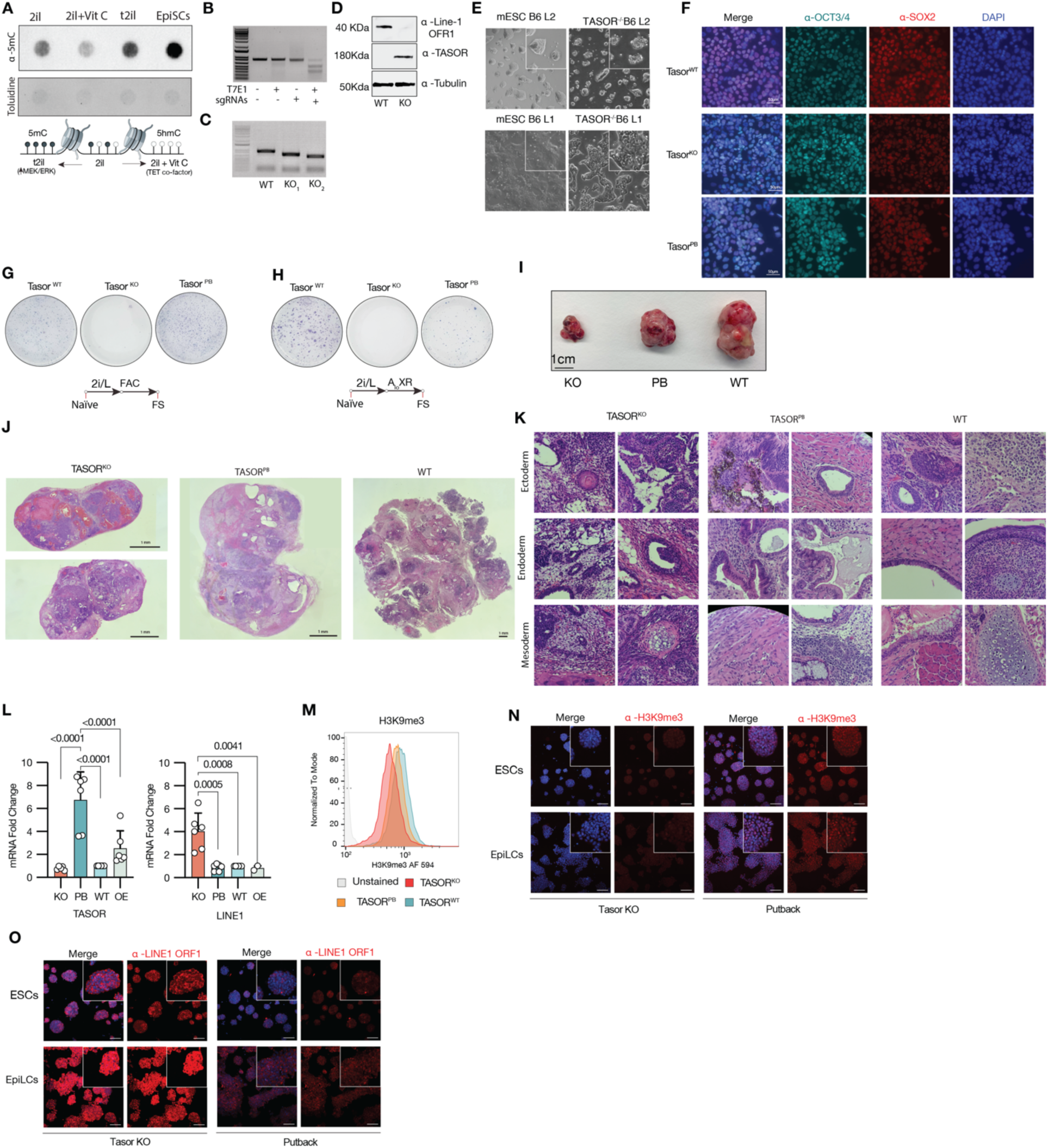
TASOR loss characterization in naïve ESCs. (**A**) DNA dot blot for 5mC and toluidine blue staining of genomic DNA from mouse ESCs cultured in 2i/L, PD03 titrated 2i/L (t2i/L), 2i/L plus vitamin C (2i/L +Vit C). (**B**) Agarose gel electrophoresis of T7 endonuclease assay for *Tasor* CRISRPR-CAS9 sgRNAS validation. (**C**) Agarose gel electrophoresis of *Tasor* knockout (KO) genotyping for line**(E)** Brightfield image of TASOR KO lines 2 and line 2. s 1 and 2. (**D**) Western blot for L1 ORF1 and TASOR in *Tasor* KO clone. (**F**) Immunofluorescence staining of OCT4, SOX2 and DAPI, for *Tasor* WT, KO, and PB cells. (**G**) Brightfield color images of colonies after FAC (also known as FTW) formative cell conversion. (**L**) Brightfield color images of colonies after AloXR formative cell conversion. (**I**) Representative teratomas of *Tasor* KO, PB, and WT naïve mPSCs. (**J**) Panoramic stich of brightfield images of hematoxylin and eosin (H&E) depicting gross tissue morphology of *Tasor* KO, PB and Wild type teratomas. (**K**) H&E staining of *Tasor* KO, PB and Wild type teratomas depicting histological morphology for Ectoderm, Endoderm, and Mesoderm. (**L**) RT-qPCR results for TASOR and L1 in KO, PB, WT and OE. (**M**) Flow cytometry analysis for global levels of H3K9me3 in *Tasor* KO, PB, WT and OE naïve cells. (**N**) Immunofluorescent staining for H3K9me3 in *Tasor* Ko and Putback naïve mPSCs (2i/L) and EpiLCs (FA). (**O**) Immunofluorescent staining for L1 ORF1 in *Tasor* Ko and Putback naïve mPSCs (2i/L) and EpiLCs (FA).

**Supplementary Figure 2.**
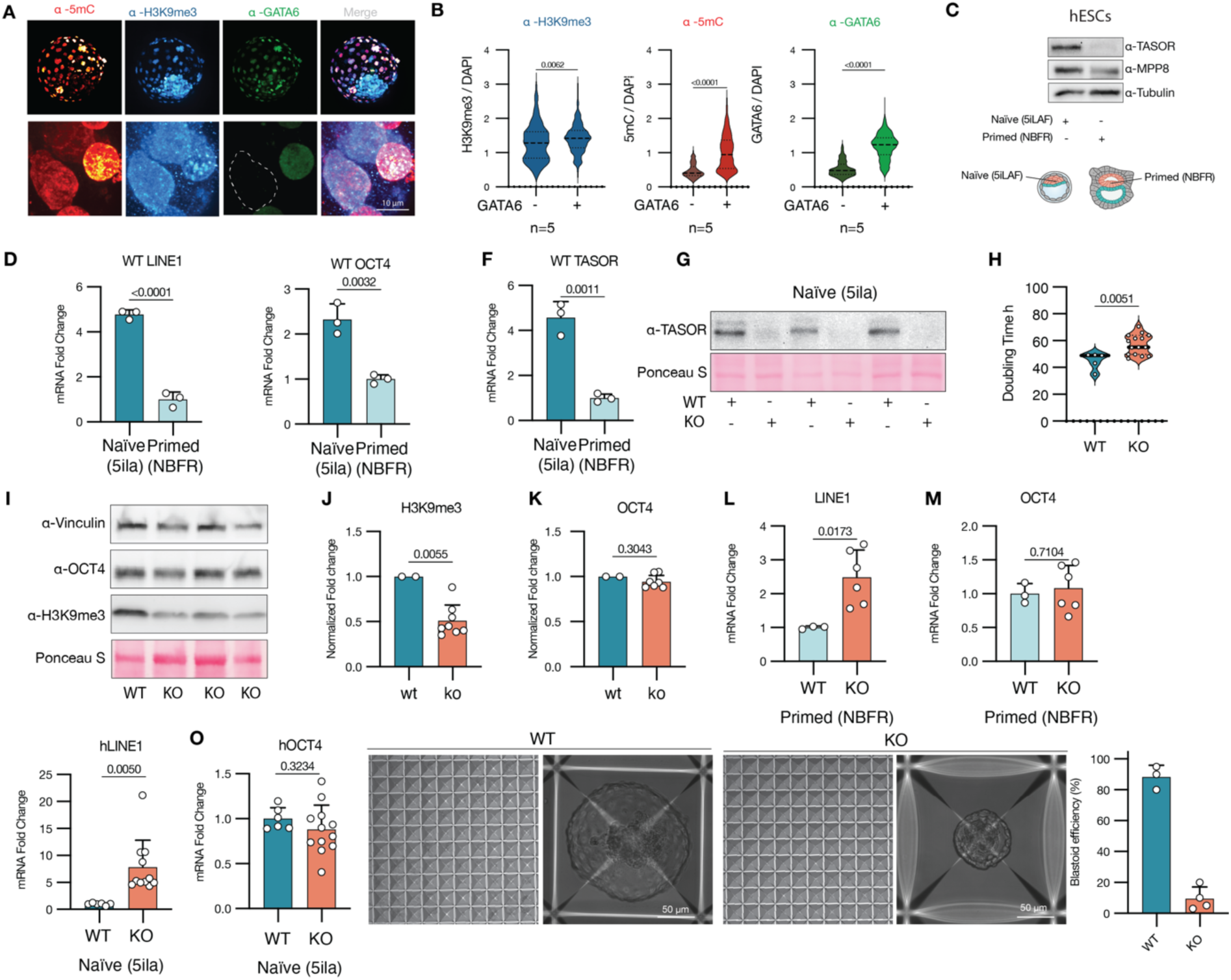
TASOR loss characterization in human PSCs. (**A**) Representative immunofluorescent staining of human blastoids derived from naïve human ESCs for differentiation marker GATA6, H3K9me3, and DNA 5mC methylation. (**B**) Quantification of H3K9me3, 5mC and GATA6, comparing GATA6 positive (Trophectoderm and hypoblast) and GATA6 negative (Epiblast) cells in Wild type blastoids, depicticing 5mC methylation gain upon exit of naïve pluripotency. (**C**) Western blot of human ESCs cultured in 5i/la (Naïve) and ESCs cultured in NBFR (Primed). RT-qPCR mRNA fold change between naïve (5ila) and primed (NBFR) human WIBR3 ΔPE-OCT4 GFP ESCs for (**D**) L1 (L1P), (**E**) OCT4, and (**F**) TASOR. (**G**) Western blot for *TASOR* knockout clones in human naïve WIBR3 ΔPE-OCT4 GFP ESCs. (**H**) Doubling time between *Tasor* Wild type (WT) and knockout (KO) cells. (**I**) Western blot for *Tasor* KO and WT clones in human naïve WIBR3 ΔPE-OCT4 GFP cells, for OCT4, and H3K9me3. (**J**) Loading control normalized relative protein level between TASOR KO and WT for H3K9me3. (**K**) Loading control normalized relative protein level between KO and WT for OCT4 (**L**) RT-qPCR mRNA fold change between WT and KO primed human ESCs for L1 (L1P). (**M**) RT-qPCR mRNA fold change between WT and KO primed human ESCs for OCT4. (**N**) RT-qPCR mRNA fold change between WT and KO naïve human ESCs for L1. (**O**) RT-qPCR mRNA fold change between WT and KO naïve human ESCs for OCT4. (**P**) Brightfield image of aggrewell plate depicting blastoid formation efficiencies in TASOR WT and KO ESCs. (**Q**) Quantification of blastoid formation efficiency between TASOR WT and KO ESCs.

**Supplementary Figure 3.**
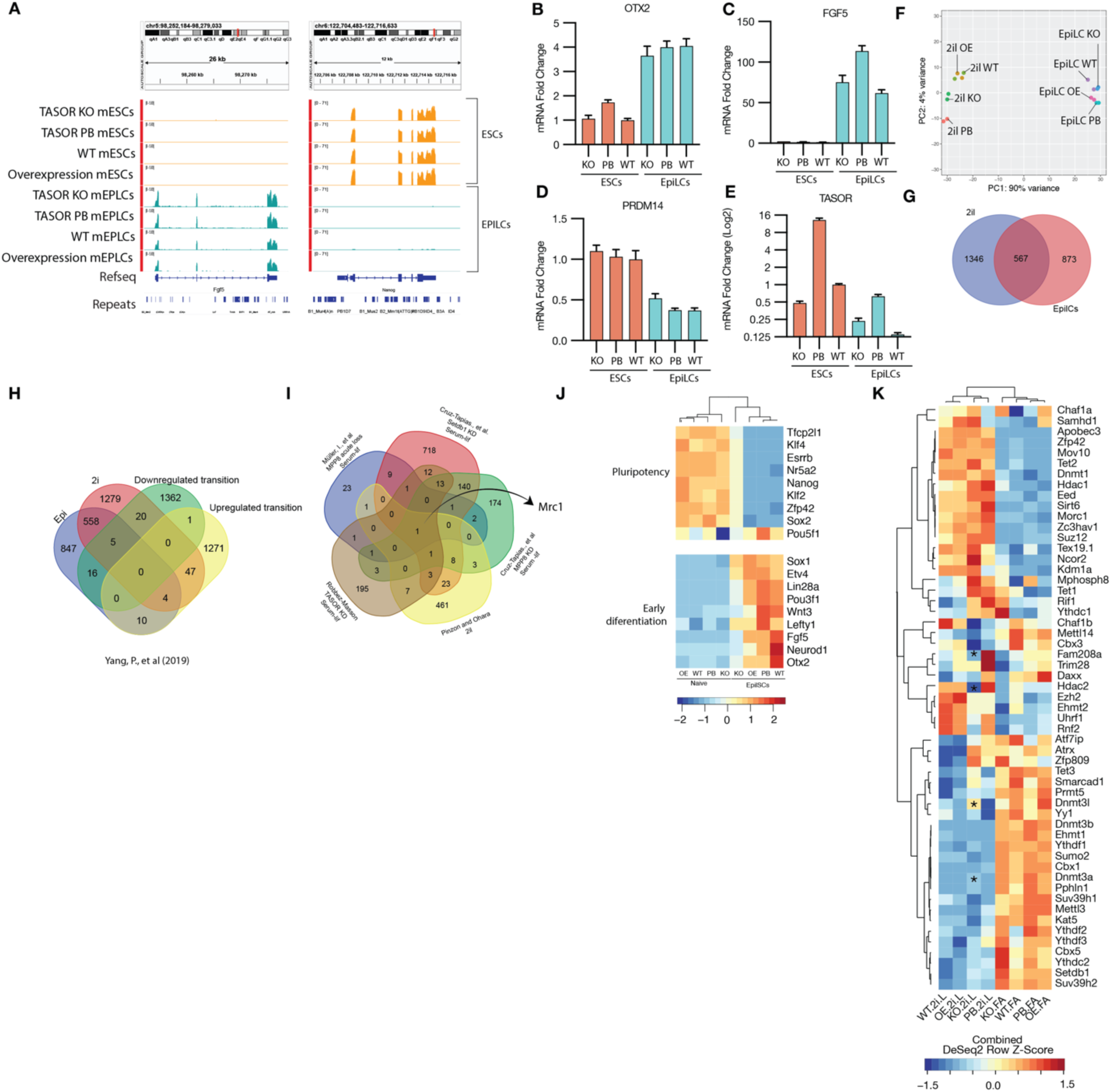
RNAseq characterization of naïve ESCs and EpiLCs upon TASOR loss. (**A**) IGV RNAseq tracks for *Tasor* knockout (KO), putback (PB), and Wild type (WT), overexpression (OE) naïve mouse ESC (2i/L) and EpiLCs (FA) depicting expression of FGF5 and NANOG. (**B**) RT-qPCR mRNA fold change between WT, KO and PB mouse ESCs and EpiLCs for *OTX2*. (**C**) RT-qPCR mRNA fold change between WT, KO and PB mouse ESCs and EpiLCs for *FGF5*. (**D**) RT-qPCR mRNA fold change between WT, KO and PB mouse ESCs and EpiLCs for *PRDM14*. (**E**) RT-qPCR mRNA fold change between WT, KO and PB mouse ESCs and EpiLCs for *Tasor*. (**F**) Principal component analysis of Naïve and EpiLCs samples. (**G**) Venn diagram of differentially expressed genes (DEGs) between TASOR KO and WT, PB, OE in naïve and EpiLCs. (**H**) Venn diagram of DEGs in naïve ESCs (2i/L) and EpiLCs (48h FA) compared to up regulated and downregulated genes in Naïve to EpiLCs trasntion in the in the stem cell atlas dabase^22^. (**I**) Venn diagram shown the common upregulated genes between our datasets and Cruz-Tapias., et al MPP8 knockdown and SETDB1 knockdown ^106^, Müller, I., et al MPPP8 acute loss^54^ and Robbez-Masson^18^. (**J**) Heatmap of different pluripotency and early differentiation markers between *Tasor* KO, PB, WT, OE naïve mouse ESC (2i/L) and EpiLCs (FA). (**K**) Heatmap of different epigenetic markers between *Tasor* KO, PB, WT, OE naïve mouse ESC (2i/L) and EpiLCs (FA).

**Supplementary Figure 4.**
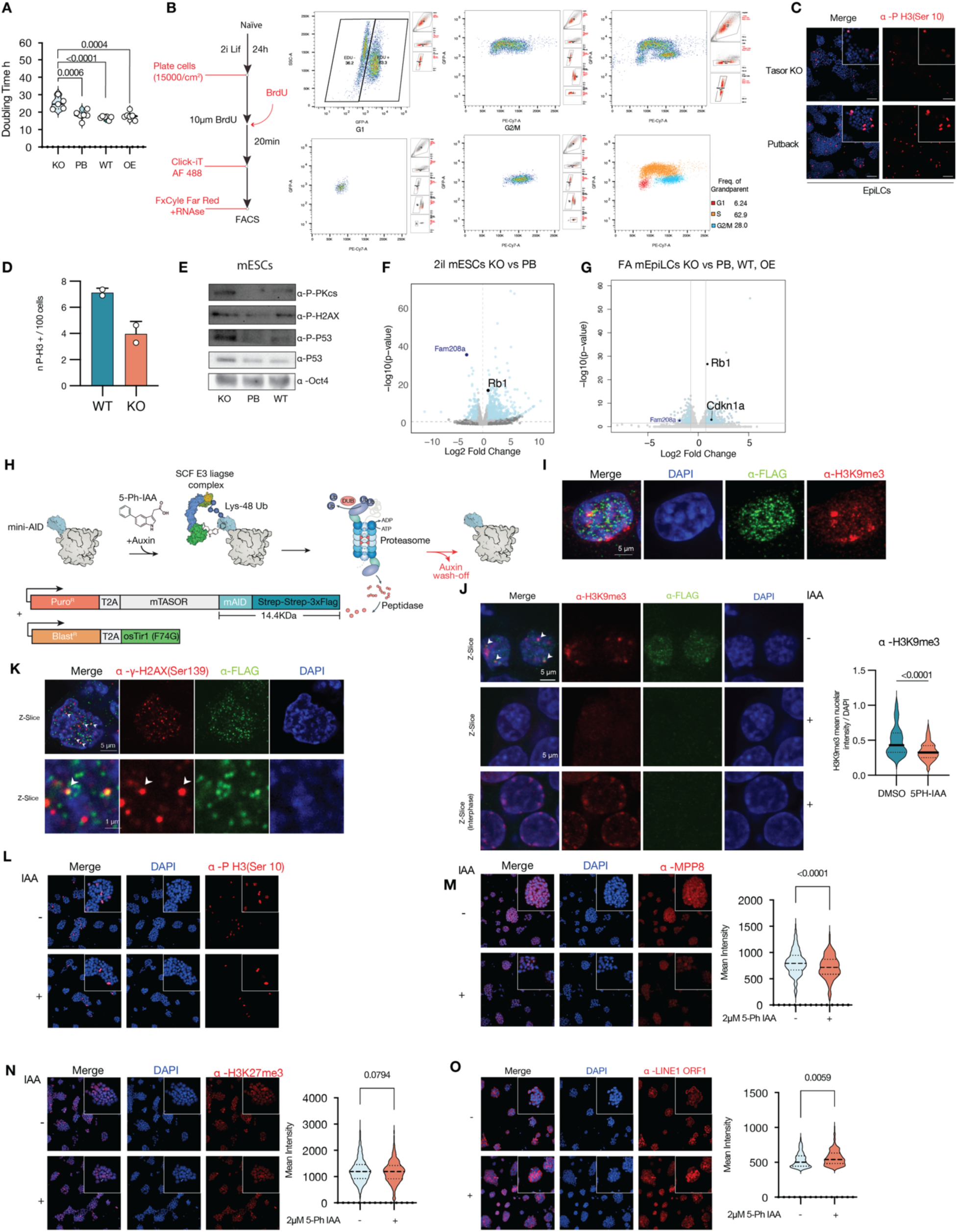
*TASOR* loss induces DNA damage and cell cycle arrest. (**A**) Doubling time of *Tasor* KO, putback (PB), Wild type (WT) and overexpression (OE). (**B**) Diagram for cell cycle analysis via BrdU incorporation and Click-it AF488 staining with FxCycle far red DNA stain with RNAse treatment, gating strategy to separate S, G1 and G2/M population is shown, percentages of grandparent population are shown. (**C**) Immunofluorescence staining for for the mitosis marker phospho H3 (Serine 10) of KO EpiLCs (FA). (**D**) Quantification of phospho H3 positive cells per 100 cells. (**E**) Western blot for DNA damage markers phospho DNA protein kinase catalytic subunit (DNAPKcs) serine 2056, for phosphoserine 139 of histone H2AX (γH2AX), and phosphoserine 15 of P-53 in naïve cells. (**F**) Volcano plot for naïve ESCs compared to putback, depicting the upregulation of *RB1*transcript. (**G**) Volcano plot for EpiLCs compared to PB, WT and OE, depicting the derepression of *Rb1* and Cdkn1a (*P21*). (**H**) Diagram of the auxin-inducible degron 2 constructs. (**I**) Z-slice confocal immunofluorescence image for H3K9me3 and FLAG; white arrows indicate colocalization spots. (**J**) Z-Slice confocal image of immunofluorescence staining for H3K9me3 and Flag with DAPI counterstaining for TASOR-mAID-TAP, bottom depicts cells in interphase with H3K9me3 chromocenters associated to the nuclear lamina. (**K**) Z-Slice confocal image of immunofluorescence staining for phosphoserine 139 of histone H2AX (γH2AX) and Flag with DAPI counterstaining for TASOR-mAID-TAP. (**L**) Maximum intensity projection immunofluorescence staining for phosphor H3 of Tasor knockout (KO) naïve mPSCs recued with TASOR-mAID-TAP with or without 2µM 5-Ph IAA during 48h for (**L**) phospho H3, (**M**) MPP8, (**N**) H3K27me3, and (**O**) L1 ORF1.

**Supplementary Figure 5.**
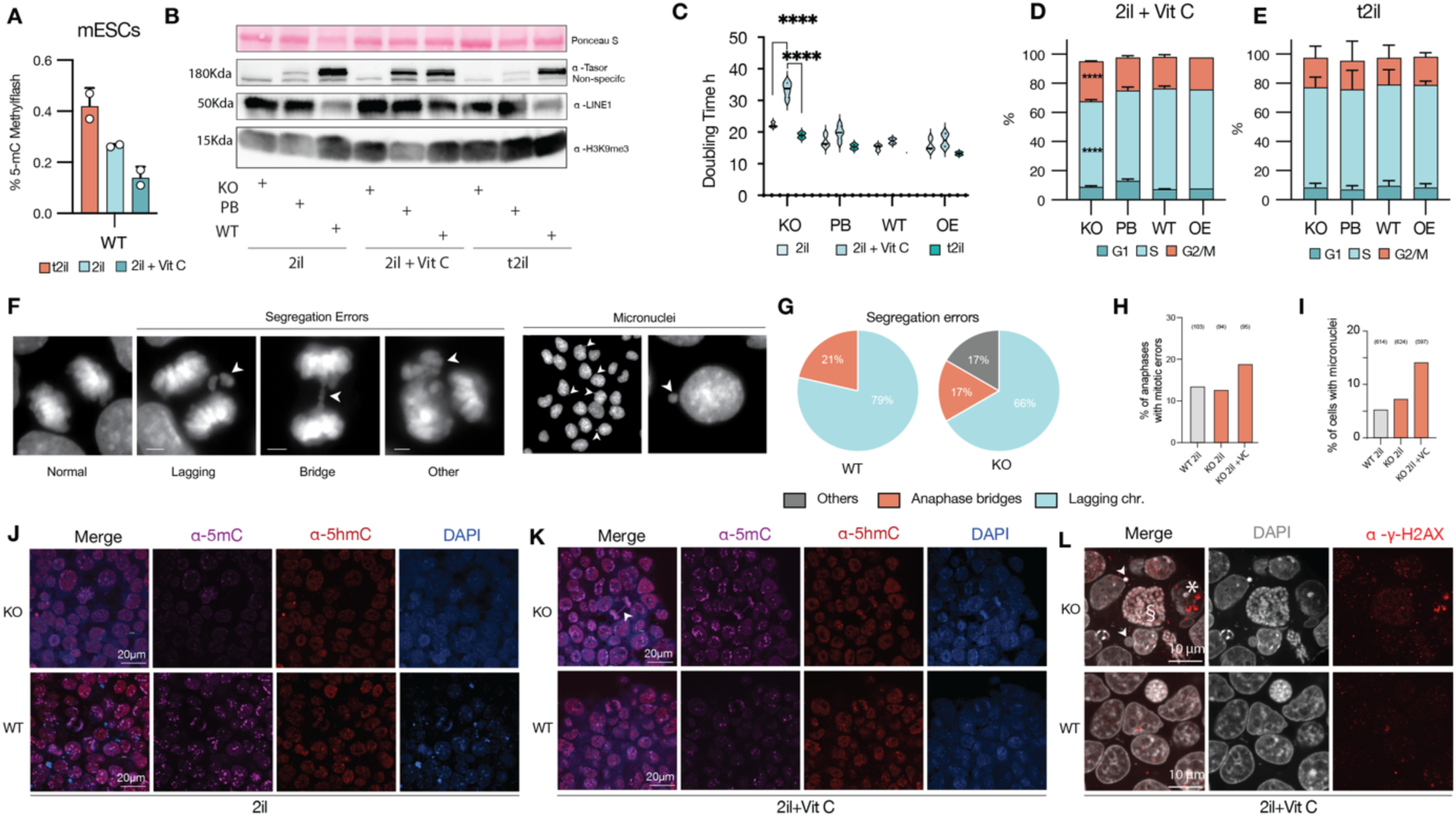
*Tasor* KO naïve mPSCs are sensitized to Vit C induced hypomethylation. (**A**) Methylflash quantification of 5mC levels in naïve mPSCs cultured in cultured in 2i/L (1µM PD03), t2i/L (0.3µM PD03) and 2i/L with vitamin C. (**B**) Flow cytometry cell cycle analysis via Edu Incorporation and DNA staining for cells in 2i/L+Vit C. (**C**) Flow cytometry cell cycle analysis via Edu Incorporation and DNA staining for cells in t2i/L. (**D**) Doubling time quantification in naïve mPSCs *Tasor* knockout (KO), putback (PB), Wild type (WT), and overexpression (OE) cultured in 2i/L, t2i/L and 2i/L with vitamin C, depicting *Tasor* KO naïve mPSCs sensitivity to vitamin C. (**E**) Western blot of *Tasor* knockout (KO), putback (PB), and Wild type (WT) in cells cultured in 2i/L, t2i/L and 2i/L with vitamin C for H3K9me3, L1 ORF-1 and TASOR. (**F**) Confocal epifluorescence images depicting different types of segregation errors and micronuclei. (**G**) Quantification of segregation erros between mouse *Tasor* knockout (KO) and Wild type (WT). (**H**) Quantification of percent of anaphases with mitotic errors between mouse *Tasor* knockout (KO) and Wild type (WT) in 2i/L or 2i/L plus vitamic C. (**I**) Quantification of percent of cells with micronuclei between mouse *Tasor* knockout (KO) and Wild type (WT) in 2i/L or 2i/L plus vitamic C. (**J**) Immunofluorescence staining for 5mC and 5hmC with DAPI counterstaining of *Tasor* knockout (KO) and wiltype (WT) naïve mPSCs cultured in 2i/L. (**K**) Immunofluorescence staining for 5mC and 5hmC with DAPI counterstaining of *Tasor* knockout (KO) and wiltype (WT) naïve mPSCs cultured in 2i/L plus vitamin C. (**L**) Z-slice confocal immunofluorescence for phosphoserine 139 of histone H2AX (γH2AX) with DAPI counterstaining of *Tasor* knockout (KO) naïve mPSCs cultured in 2i/L plus vitamin c, depicting DNA damage (*), micronuclei (arrow head) and accumulation of abnormal karyotype(§).

**Supplementary Figure 6.**
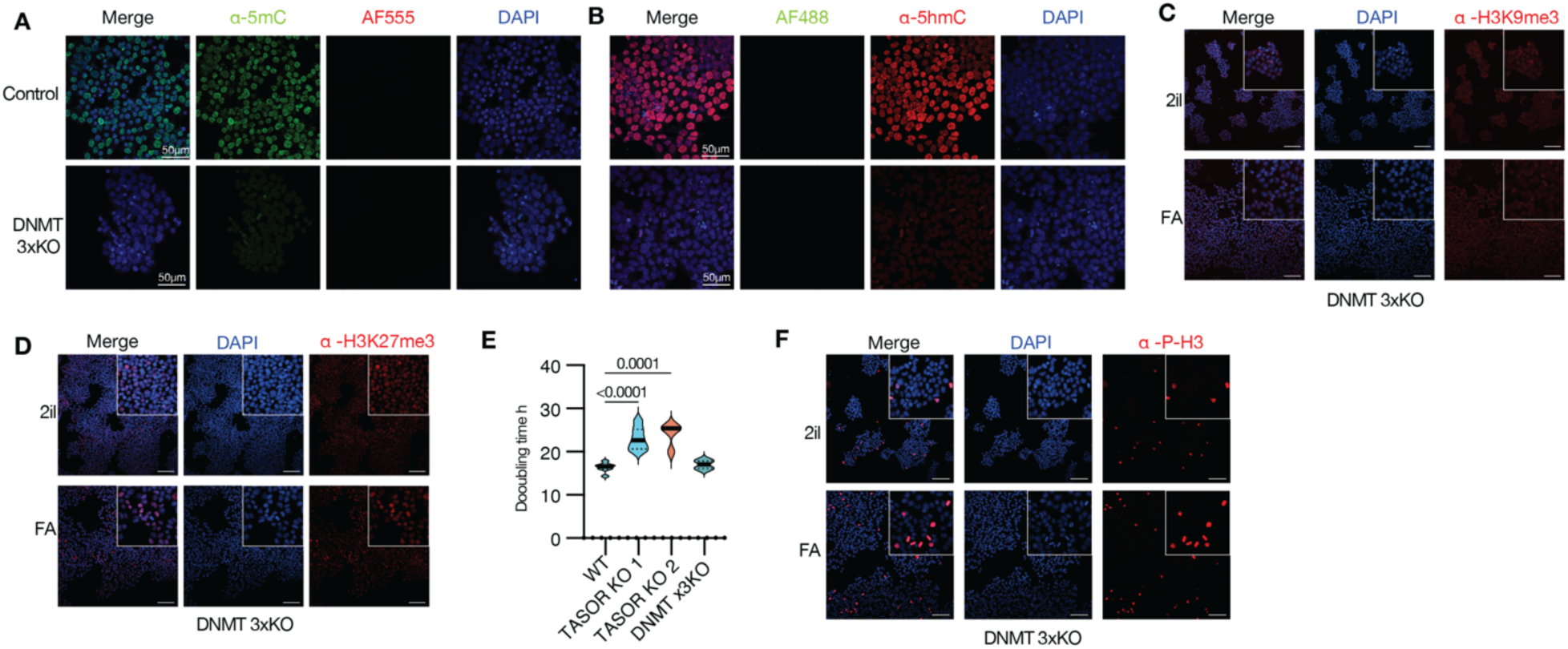
Characterization of DNMTx3KO naïve ESCs and EPiLCs. (**A**) Immunofluorescence staining for 5mC with DAPI counterstaining for Dnmt1, Dnmt3a and Dnmt3b triple knockout (Dnmt x3KO) and control naïve ESCs cultured in 2i/L. (**B**) Immunofluorescence staining for 5hmC with DAPI counterstaining for Dnmt x3KO and control naïve ESCs cultured in 2i/L. (**C**) Immunofluorescent staining for H3K9me3 in Dnmt x3KO naïve ESCs (2i/L) and EpiLCs (FA). (**D**)Immunofluorescent staining for H3K27me3 in Dnmt x3KO naïve mPSCs (2i/L) and EpiLCs (FA). (**E**)Doubling time of Wild type, *Tasor* KO and Dnmt x3KO naïve mPSCs cultured in 2i/L. (**F**) Immunofluorescence staining for for the mitosis marker phospho H3 (Serine 10) of Dnmt triple KO naïve mPSCs (2i/L) and EpiLCs (FA).

**Supplementary Figure 7.**
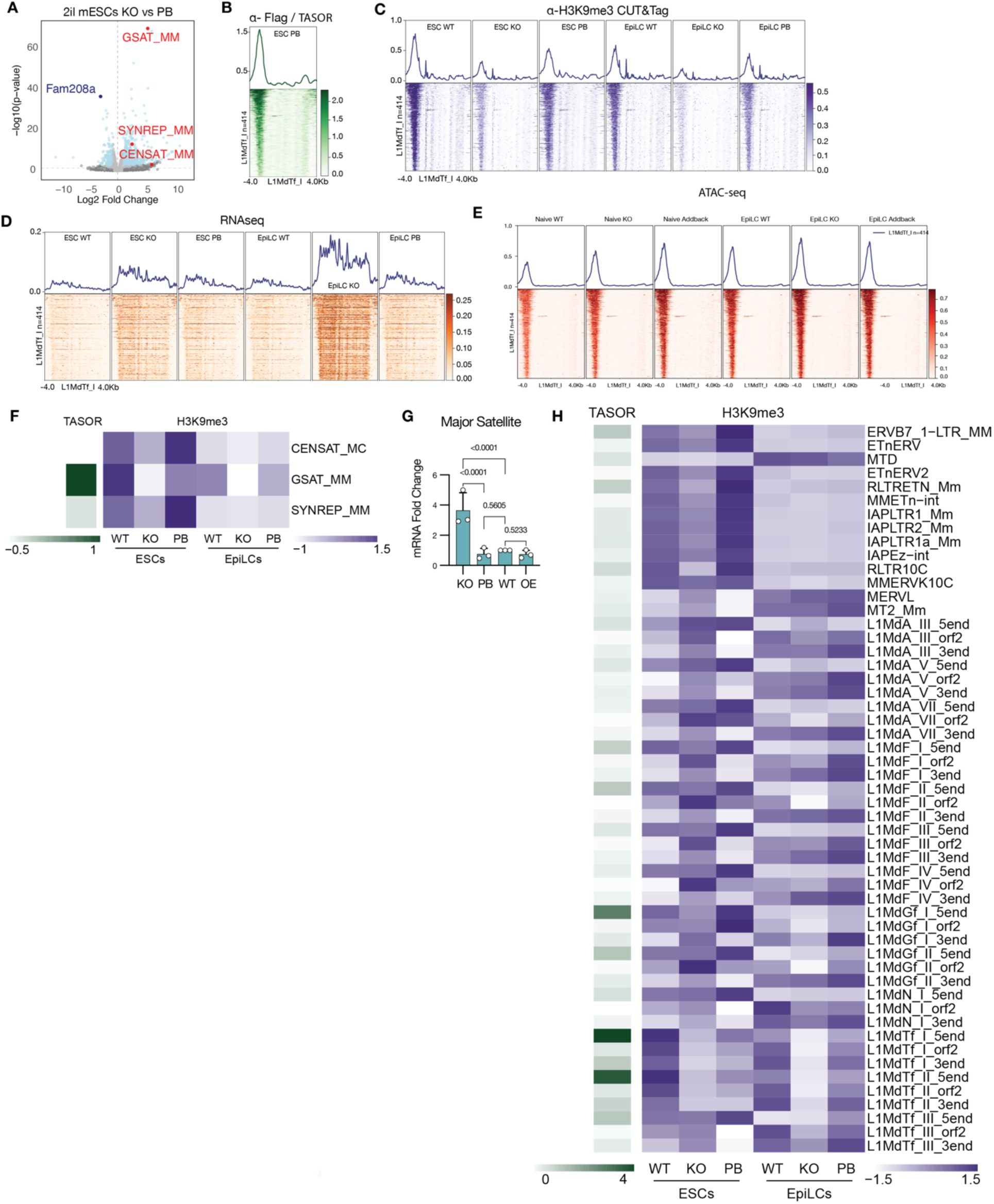
Epigenetic profiling of H3K9me3 and TASOR-3xFLag reveals Tasor regulation of L1 and repeats. (**A**) Volcano plot for naïve ESCs compared to putback, depicting the derepression of General satellite repeats (GSAT_MM), Simple or minor Repeats (SYNREP_MM), and Centromeric satellite repeats (CENSAT_MC). (**B**) Average CUT&Tag profiles (top) and heatmaps (bottom) at L1MdTf_I (n=414) for TASOR-3xFlag. (**C**) Average CUT&Tag profiles (top) and heatmaps (bottom) at L1MdTf_I (n=414) for H3K9me3. (**D**) Average RNAseq profiles (top) and heatmaps (bottom) at L1MdTf_I (n=414). (**E**) Average ATACseq profiles (top) and heatmaps (bottom) at L1MdTf_I (n=414) for H3K9me3. (**F**) Heatmap for CUT&Tag of TASOR-3xFlag and H3K9me3 at General satellite repeats (GSAT_MM), Simple or minor Repeats (SYNREP_MM), and Centromeric satellite repeats (CENSAT_MC). (**G**) RT-qPCR mRNA fold change between *Tasor* KO, PB, WT naïve mouse ESC (2i/L) for Major satellite repeats RNA. (**H**) Heatmap for CUT&Tag of TASOR-3xFlag and H3K9me3 at different retroviral families.

**Supplementary Figure 8.**
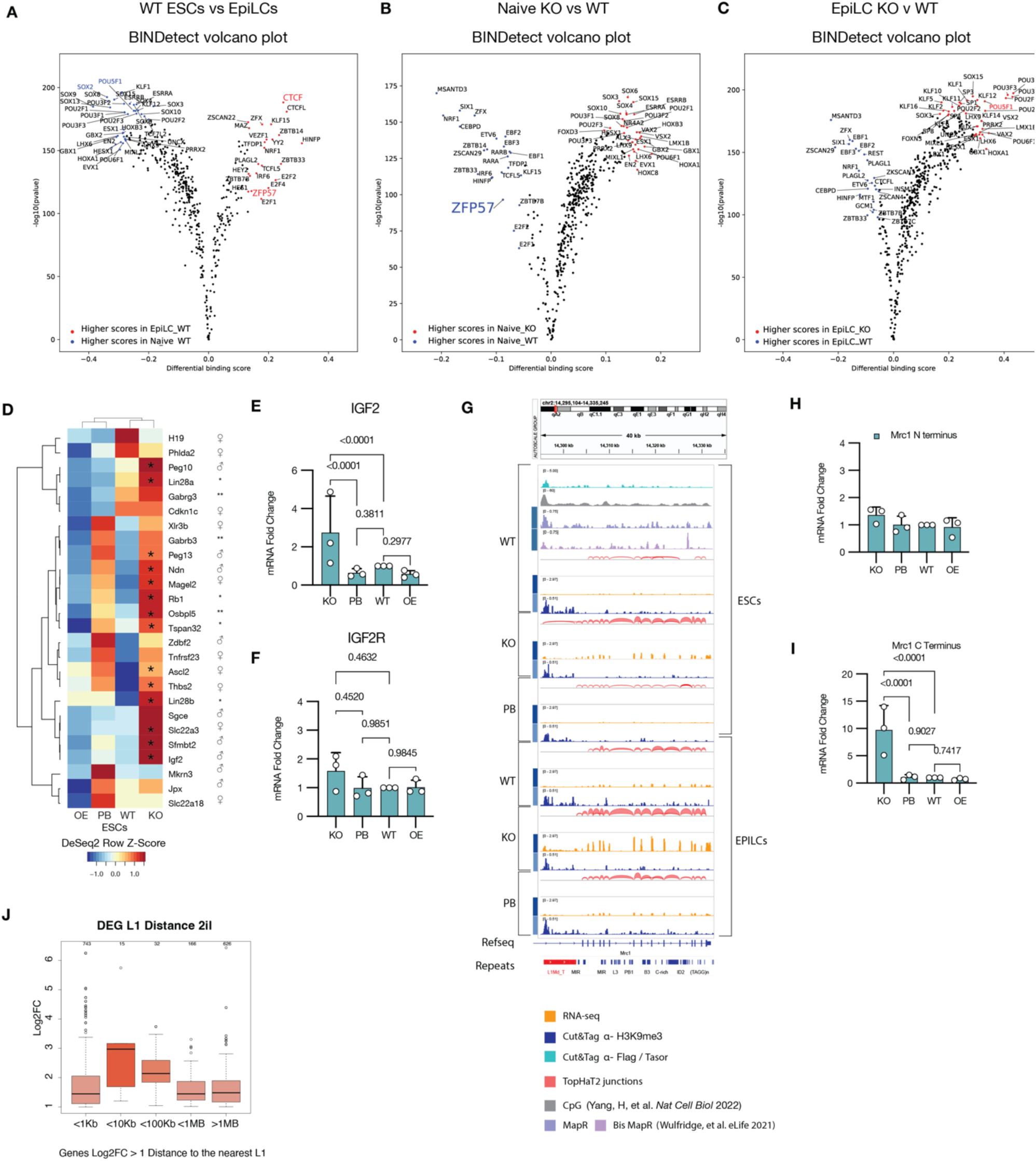
Analysis of Tasor KO cells differentially expressed genes and ATAC peaks. (**A**) Differential transcription factor binding scores from ATAC data using BINDetect between Wild type naïve mouse ESC (2i/L) and EpiLCs (FA). (**B**) Differential transcription factor binding scores from ATAC data using BINDetect between *Tasor* KO versus WT naïve mouse ESC (2i/L). (**C**) Differential transcription factor binding scores from ATAC data using BINDetect between *Tasor* KO versus WT EpiLCs. (**D**) Heatmap of different differentially expressed imprinted genes in naïve mouse ESC (2i/L). * Represent significant differentially expressed genes, log2 fold change ≥1, pval≤0.05. (**E**) RT-qPCR mRNA fold change between *Tasor* KO, PB, WT naïve mouse ESC (2i/L) for *IGF2*. (G) (**F**) RT-qPCR mRNA fold change between *Tasor* KO, PB, WT naïve mouse ESC (2i/L) for IGF2 receptor *(IGF2R)*. (**G**) IGV tracks for RNAseq (Orange), Cut&Tag tracks for TASOR-3xFLAG(Teal), H3K9me3 (dark blue), ToPHAT2 junctions(red), CpG number from Yang, et al ^107^ and R-loops from Wulfridge, et al^85^, in *Tasor* KO, PB, WT naïve mouse ESC (2i/L) and EpiLCs (FA), depicting the position effect varigenation of L1s affecting nearby gene expression upon TASOR loss. and R-loops from Wulfridge, et al^85^, in *Tasor* KO, PB, WT naïve mouse ESC (2i/L) and EpiLCs (FA), depicting the position effect varigenation of L1s affecting nearby gene expression upon TASOR loss. (**H**) RT-qPCR mRNA fold change between *Tasor* KO, PB, WT naïve mouse ESC (2i/L) for *Mrc1* N terminus. (**I**) RT-qPCR mRNA fold change between *Tasor* KO, PB, WT naïve mouse ESC (2i/L) for *Mrc1* C terminus. (**J**) Upregulated DEG genes nearest Line 1 distance in 2i/L.

**Supplementary Figure 9.**
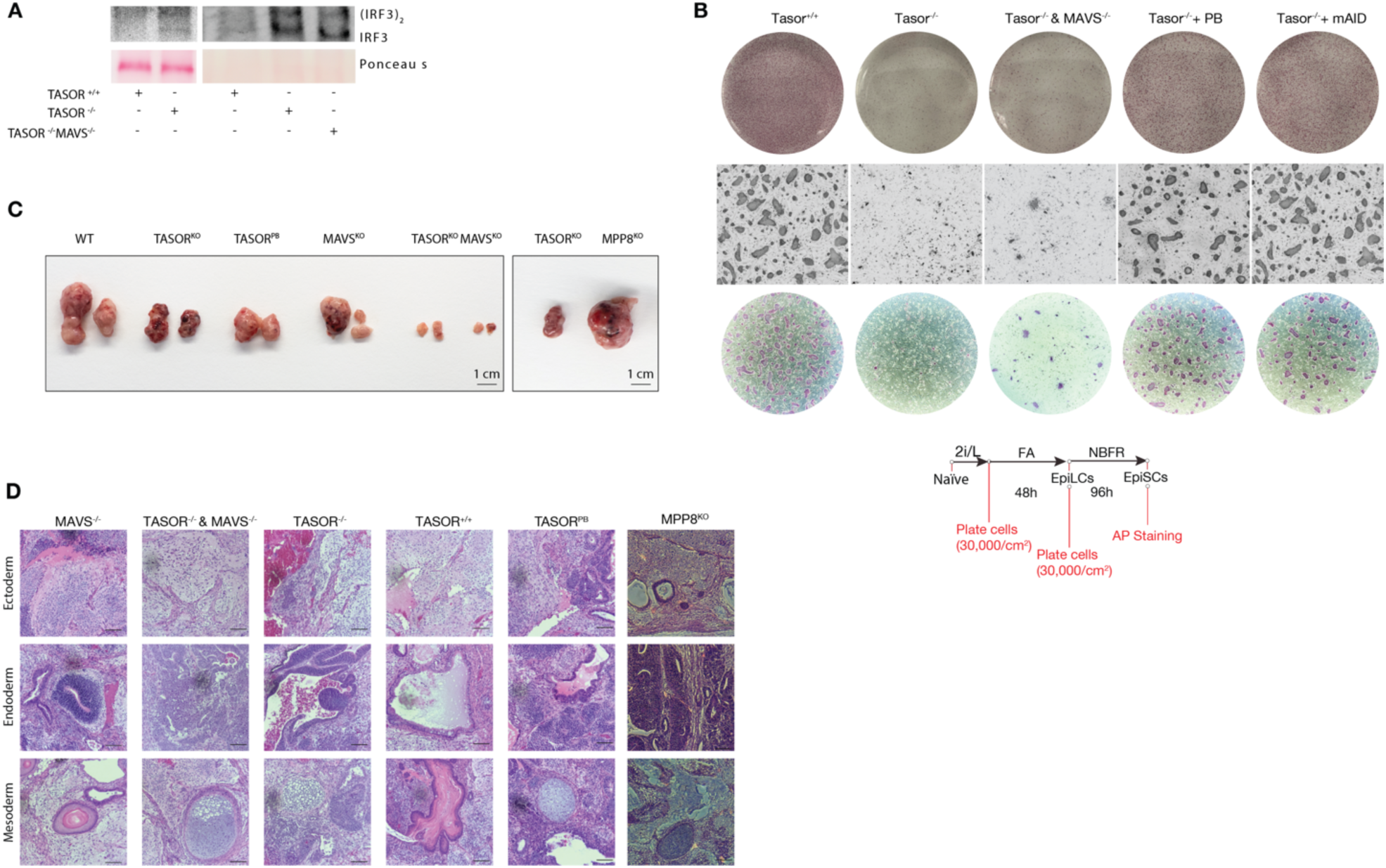
Characterization of MAVS and TASOR double Knockout cells. (**A**) Westren blot of native PAGE gel for IRF3 dimerization in *Tasor* KO, WT or *Tasor* and *MAVS* double KO. (**B**) Alkaline phosphatase of colony formation assay for *Tasor* WT, KO, *Tasor* and *MAVS* double KO, PB, and KO recued with TASOR-mAID-TAP naïve ESCs transitioned to EpiSCs. (**C**) Teratomas of *Tasor WT,* KO, PB, *MAVS KO, Tasor* and *MAVS* double KO, and *MPP8* KO naïve ESCs. (**D**) Brightfield histopathological slides stained with hematoxylin and eosin (H&E) of teratomas. Scale bar: 100µm.

## METHODS

### Culture of mouse embryonic stem cells (ESC)

All cell cultures were performed in N2B27 basal media. This media was prepared using the following (250ml): 125 ml DMEM/F12 (Invitrogen), 125 ml neurobasal medium (Invitrogen), 1x N2 supplement (5ml, Invitrogen), 1x B27 supplement (10ml, Invitrogen), 1× GlutaMAX (2.5ml, Gibco), 1× nonessential amino acids (2.5ml, Gibco), 0.1 mM β-mercaptoethanol (Gibco), 1% fatty acid free BSA, and 2.5µg/ml of prophylactic plasmocin (InvivoGen) to prevent mycoplasma contamination during maintenance of the cells but was removed before any experimental assay (InvivoGen).

All cell culture was performed as following: cells were washed with 1xPBS and dissociated with TrypLE (Thermo Fisher) for 3 minutes at 37°C; cells were then collected with 0.05% BSA in DMEM-F12 (Thermo Fisher) and centrifuged at 1000xg for 3 minutes and resuspended in 1ml of media per 9.6cm^2^. Each passage cells were counted using Countess II (Thermo Fisher) and plated at a density of 15,000 cells/cm^2^, at this plating ratio cells were passaged every 4 days. Fresh media (2ml per 9.6 cm^2^) was added during the first two days, and on days 3 and 4 the media amount was doubled. Cell cultures were maintained in a 37°C humidified in incubator with 5% CO_2_. Cells were cryopreserved in CryoStor CS10 (Biolife solutions) at 0.5×10^6 cells per ml in −80°C, using in CoolCell freezing containers (Corning). The following day, cells were moved for long term storage in liquid nitrogen.

Naïve mouse ESCs (naïve ESCs) were cultured feeder free, on Geltrex (Gibco) coated cell culture dishes in 2i/L media. 2i/L media was prepared in N2B27 basal media with the addition of the following small molecules and cytokines: 1 μM PD0325901 (MEK 1/2 inhibitor, Selleckchem), 3µM CHIR 99021 (WNT agonist via GSK3 α/β inhibiton, Selleckchem) and in the presence of the STAT3 agonist, leukemia inhibitory factor (LIF, 10ng/ml).

### Culture of epiblast like stem cells (EpiLCs), epiblast stem cells (EpiSCs), and colony clonogenicity formation assays

Mouse epiblast like stem cells (EpiLCs), were transitioned from 2i/L cultures by plating the cells on Geltrex (Gibco) coated cell culture dishes at a density of 15,000 cells/cm^2^ in N2B27 based FA media containing 12ng/ml of FGF2 and 20ng/ml of Activin A for 48 hours. Media was replaced every 24 hours. Mouse epiblast stem cells (EpiSCs) were transitioned from EpiLCs by plating the cells on mitotically inactivated mouse embryonic fibroblast (MEF) on 0.1% gelatin coated cell culture dishes at a density of 15,000 cells/cm^2^ in N2B27-based NBFR media containing 20ng/ml of FGF2 and 2.5µM of the WNT antagonist IWR1 (Selleckchem). Media was replaced every 24 hours. For colony clonogenicity assays, cells were plated at 500-5000 cells/cm^2^. Once colonies had grown (6-15 days), cells were stained with an alkaline phosphatase staining kit following manufactures instructions (Abcam), or with coomassie brilliant blue (Thermo)

### Culturing of human naïve PSCs

Naïve WIBR3 human ES cells were obtained from R. Jaenisch and T. Theunissen. Primed human PSC were cultured in MEFs on 0.1% gelatin-coated cell culture dishes on N2B27 based media as described above. Primed cells were cultured on irradiated mouse embryonic fibroblasts in NBFR (Table 1) supplemented with 20ng/ml of LIF and 20ng/ml of Activin A. Primed cell lines were reset to the naïve state following a previously described protocol^4108^. Briefly, 20,000 cells/cm^2^ primed cells were treated with 1µM PD0325901(Selleckchem), 1mM Valporic acid (VPA, Medchemexpress), 20ng/ml of leukemia inhibitory factor (LIF, Peprotech) and CEPT cocktail [50 nM Chroman 1 (MedChem Express), 5 µM Emricasan (Selleckchem), 1X polyamine supplement (Sigma), and 0.7 µM TransISRIB (Tocris)], for 3 days. Then, media was changed to a modified 5i/L/A medium.

**Table 1.** Media formulations.

| **MEDIA FORMULATIONS** | **Stage** | **SOURCE** | **IDENTIFIER** |
| --- | --- | --- | --- |
| **2i/L:** N2B27 base, 10ng/ml LIF, 3μM Chir99021, 1μM PD032590. | Naïve mouse PSCs | Qi-long, Y. *et al.*, 2008, *Nature*<sup>2</sup><br>Sato, N. *et al.*, 2004. *Nat Med*<sup>5</sup> | https://doi.org/10.1038/nature06968<br>https://doi.org/10.1038/nm979 |
| **2i/L + VitC:** N2B27 base, 10ng/ml LIF, 3μM Chir99021, 1μM PD0325901, 100μg/ml L-ascorbic acid. | Naïve mouse PSCs | Blaschke, K. *et al.*, 2013, *Nature*<sup>49</sup><br>Walter, M. *et al.* 2016, *Elife*<sup>50</sup> | https://doi.org/10.7554/eLife.11418<br>https://doi.org/10.1038/nature12362 |
| **t2i/L:** N2B27 base, 10ng/ml LIF, 3μM Chir99021, 0.2μM PD0325901, 1% FBS. | Naïve-like mouse PSCs | Gretarsson, K.H, *et al.*, 2020, *Nat Struct Mol Biol*<sup>43</sup> | https://doi.org/10.1016/j.stemcr.2017.05.014 |
| **FA (48h):** N2B27 base, 12ng/ml FGF2 and 20ng/ml Activin A. | Epiblast-like stem cells (EpiLSCs). | Hayashi, K. *et al.*, 2011, *Cell*<sup>7</sup> | https://doi.org/10.1016/j.cell.2011.06.052 |
| **NBFR:** N2B27 base, 20 ng/mL FGF2 and 2.5 μM IWR-1. | Epiblast stem cells (EpiSCs). | Brons, I. *et al.*, 200, *Nature*.<sup>8</sup><br>Tesar, P. J. *et al.*, 2007, *Nature*.<sup>9</sup><br>Wu, J. *et al.*, 2015, *Nature*<sup>29</sup> | https://doi.org/10.1038/nature05950<br>https://doi.org/10.1038/nature05972<br>https://doi.org/10.1038/nature14413 |
| **AloXR:** N2B27 base, 3ng/ml of activin A, 2μM XAV939 and 1.0μM BMS493 | Formative PSCs | Kinoshita *et al.*, 2021, *Cell Stem Cell*<sup>27</sup> | https://doi.org/10.1016/j.stem.2020.11.005 |
| **FAC:** N2B27 base, 1% BSA, , 20ng/ml Activin A, 20ng/ml FGF2 ,3 μM Chir99021. | Formative PSCs | Yu, *et al.*, 2021, *Nature* | https://doi.org/10.1038/s41586-021-03356-y |
| **5i/LA:** 1 μM PD0325901, 0.5 μM IM-12, 0.5 μM SB590885, 1 μM WH-4-023, 20 ng/ml recombinant human LIF, 10 ng/ml Activin A, and 5 μM Y-27632. | Naïve human PSCs | Theunissen, T. W., *et al.* 2014, *Cell Stem Cell* | https://doi.org/10.1016/j.stem.2014.07.002 |
| **eHDM:** N2B27 base, 1% BSA, 20 ng/ml bFGF (Peprotech), 20 ng/ml activin A, 3 μM CHIR99021 and CEPT cocktail [50 nM Chroman 1 , 5 μM Emricasan, 1X polyamine supplement, and 0.7 μM TransISRIB ]. | Enhanced hypoblast differentiation media (eHDM) | L. Yu, D. Logsdon, C. A. Pinzon-Arteaga., et al, 2023, *Cell Stem Cell*<sup>108</sup> | https://doi.org/10.1016/j.stem.2023.08.002 |
| **eTDM** N2B27 base, 1% BSA, 1 μM PD0325901, 2 μM A83-01( or 1 μM A83-01, 1 μM A77-01), 0.5 μM SB590885, 1 μM WH-4-023, 10 ng/ml LIF, 0.5 μM LPA and and CEPT cocktail [50 nM Chroman 1 , 5 μM Emricasan , 1X polyamine supplement, and 0.7 μM TransISRIB]. | Enhanced throphectoderm differentiation media (eTDM) | L. Yu, D. Logsdon, C. A. Pinzon-Arteaga., et al, 2023, *Cell Stem Cell*<sup>108</sup> | https://doi.org/10.1016/j.stem.2023.08.002 |

All naïve human ESCs were cultured on Geltrex-coated dishes. Briefly, 25,000 cells/cm^2^ naïve PSCs were plated into Geltrex-coated cell culture plates in the modified 5i/L/A medium^108^. The cells were passaged as described above with 1xCEPT cocktail^109^ [50 nM Chroman 1 (MedChem Express), 5 µM Emricasan (Selleckchem), 1X polyamine supplement (Sigma), and 0.7 µM TransISRIB (Tocris)], for 12 hours. The modified 5i/L/A medium was prepared using N2B27 basal media with the addition of 0.5% KSR, 50 µg/ml of bovine serum albumin (BSA, Sigma) and the following small molecules and cytokines: 1 μM PD0325901 (Selleckchem), 0.5 μM IM-12 (Enzo), 0.5 μM SB590885 (R&D systems), 1 μM WH-4-023 (Selleckchem), 20 ng ml−1 recombinant human LIF (Peprotech), 10 ng ml−1 Activin A (Peprotech) and 5 μM Y-27632 (Selleckchem)., 2µM XAV939 (MedChem Express) and 2µM Gö 6983 (MedChem Express). Naïve ESCs were cultured for a minimum of 10 days in the modified 5i/L/A before any experiment. Naïve human ESCs were never exposed to serum.

### Generation of blastoids from human naïve PSCs

Blastoid generation was performed as described by Yu; et al^108^. 5iLA naïve human PSCs were dissociated into single cells by incubation with TrypLE (Thermo Fisher) for 3-5 min at 37°C. Cells were collected with 0.1% BSA in DMEM-F12 medium and centrifugated at 1000 rpm (approx. 200xg) for 3 min in a swing bucket centrifuge (Legend RT+, Thermo Fisher) and recovered in 5iLA medium with 1xCEPT and 10U/ml of DNase I (Thermo Fisher) and incubated at room temperature for 15min. Cells were then passed through a 20-μm cell strainer (Pluriselect). To select for viable cells and exclude dead cells and cell debris, cells were carefully layered on top of 10ml of 0.1% BSA DMEM-F12 in a 45° angle in a 15ml conical centrifuge tube and centrifuged at 300 rpm for 10 minutes. Supernatant was removed and pelleted cells were re-suspended and manually counted in a Neubauer counting chamber. Meanwhile, AggreWell-400 (STEMCELL Technologies) plate was prepared according to the manufacturer’s instructions. In brief, 500µl of anti-adherence solution (STEMCELL Technologies) was added to each well, the plate was centrifuged at 1500 x g for 5 min, and then incubated at room temperature for a minimum of 45 min. The corresponding number of cells (32 cells / microwell) were washed and resuspended in 1ml eHDM(3µM Chir99021, 10ng/ml FGF2-3G, 20ng/ml Activin A) with 1xCETP and seeded into one well of a precoated AggreWell-400 24-well plate in 1ml of media. Each well was carefully mixed using a P200 pipette and the plate was left alone for 15 minutes to ensure equal distribution of the cells inside the well. The plate was centrifuged at 1000 x g for 3 min and cultured at 37°C in 5% CO_2_ and 5% O_2_. The day of cell plating was designated as day 0. eHDM was completely changed to eTDM on day 1 by carefully removing as much eHDM as possible without disturbing the aggregates by tilting the plate in a 45° angle, each well was washed once with 200µl of eTDM, finally 1 ml of eTDM was slowly added 200µl at a time. On the remaining days, fresh eTDM with fresh LPA was half changed every day. The human blastoids usually formed after four days of culture in eTDM. All blastoids were manually isolated using a mouth pipette under a stereomicroscope for downstream experiments. For eHDM and eTDM recepie see Table 1.

### Immunofluorescent staining

Samples (cells, and blastoids) were fixed with 4% paraformaldehyde (PFA) in 1xDPBS with 0.1% PVA for 20 min at room temperature, washed in wash buffer (0.1% Triton X-100, 5% BSA in 1xDPBS) for 15 minutes and permeabilized with 0.1-1% Triton X-100 in PBS for 1 h. For 5mC / 5hmC staining samples were treated with 4N HCL for 15 minutes, and then neutralized with 100mM Tris-HCL pH 8.0 for 30 minutes. Samples were washed 3 times in wash buffer for 5 minutes and then blocked with blocking buffer (PBS containing 5% Donkey serum, 5% BSA, and 0.1% Triton X-100) at room temperature for 1 h, or overnight at 4°C.

Primary antibodies were diluted in blocking buffer (Table 2). Blastoids were incubated in primary antibodies in U bottom 96 well plate for 2 h at room temperature or overnight at 4°C. Samples were washed three times for 15 minutes with wash buffer, and incubated with fluorescent –dye-conjugated secondary antibodies (AF-488, AF-555 or AF-647, Invitrogen) diluted in blocking buffer (1:300 dilution) for 2hr at room temperature or overnight at 4°C. Samples were washed three times with wash buffer. Finally, cells were counterstained with 300 nM 4′,6-diamidino-2-phenylindole (DAPI) solution at room temperature for 20 min.

**Table 2.**
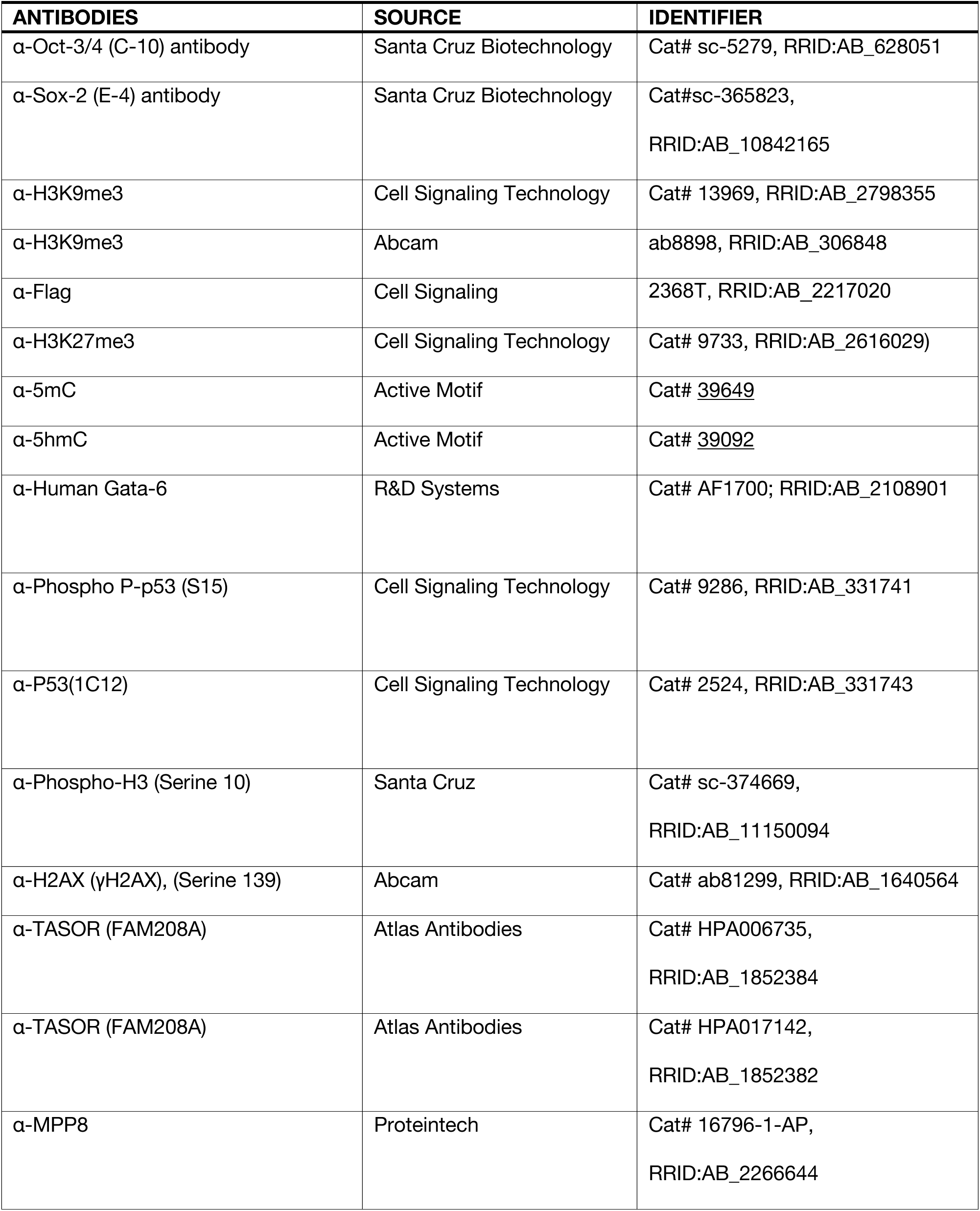
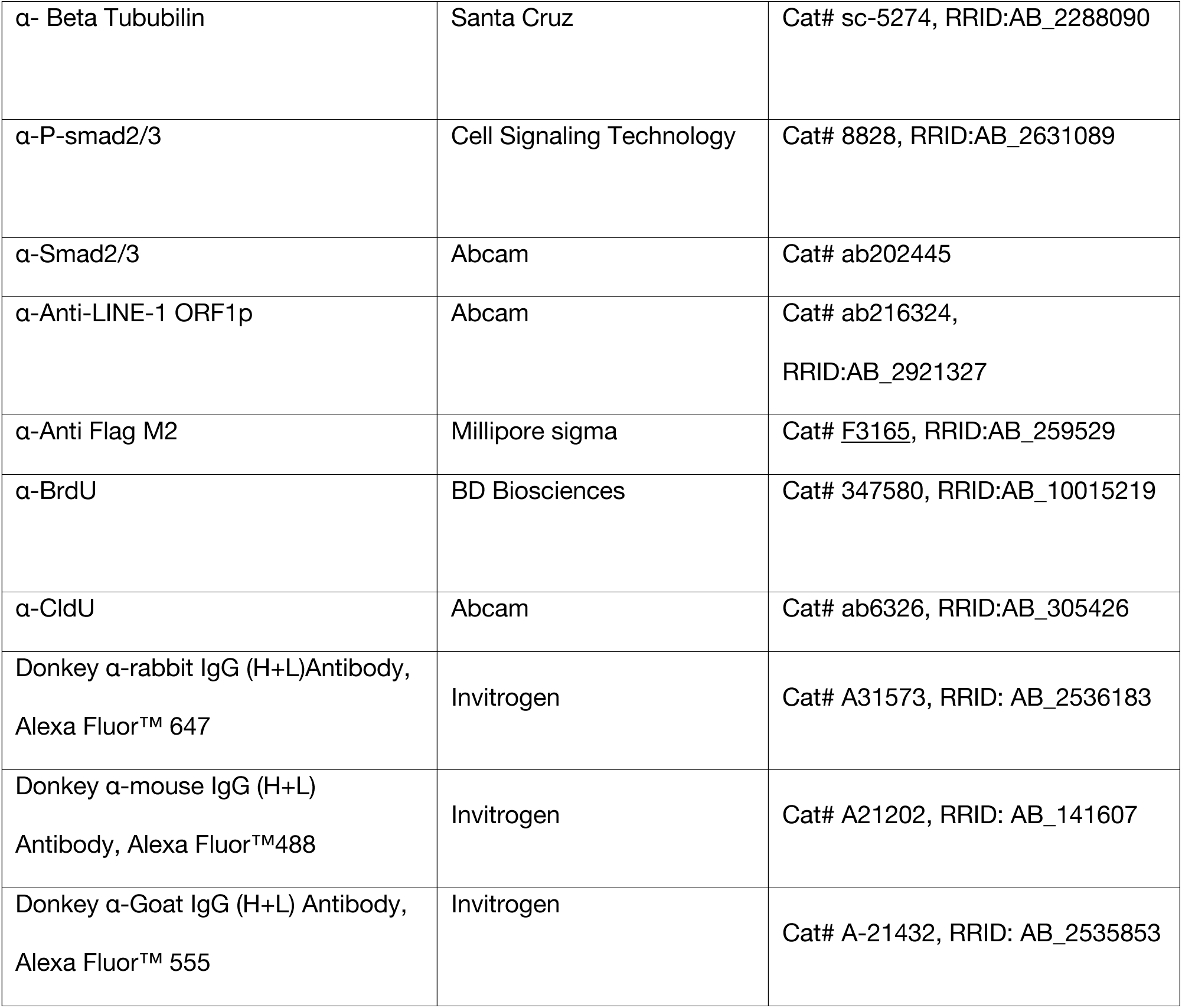
Antibodies.

| **ANTIBODIES** | **SOURCE** | **IDENTIFIER** |
| --- | --- | --- |
| α-Oct-3/4 (C-10) antibody | Santa Cruz Biotechnology | Cat# sc-5279, RRID:AB_628051 |
| α-Sox-2 (E-4) antibody | Santa Cruz Biotechnology | Cat#sc-365823,<br>RRID:AB_10842165 |
| α-H3K9me3 | Cell Signaling Technology | Cat# 13969, RRID:AB_2798355 |
| α-H3K9me3 | Abcam | ab8898, RRID:AB_306848 |
| α-Flag | Cell Signaling | 2368T, RRID:AB_2217020 |
| α-H3K27me3 | Cell Signaling Technology | Cat# 9733, RRID:AB_2616029) |
| α-5mC | Active Motif | Cat# 39649 |
| α-5hmC | Active Motif | Cat# 39092 |
| α-Human Gata-6 | R&D Systems | Cat# AF1700; RRID:AB_2108901 |
| α-Phospho P-p53 (S15) | Cell Signaling Technology | Cat# 9286, RRID:AB_331741 |
| α-P53(1C12) | Cell Signaling Technology | Cat# 2524, RRID:AB_331743 |
| α-Phospho-H3 (Serine 10) | Santa Cruz | Cat# sc-374669,<br>RRID:AB_11150094 |
| α-H2AX (γH2AX), (Serine 139) | Abcam | Cat# ab81299, RRID:AB_1640564 |
| α-TASOR (FAM208A) | Atlas Antibodies | Cat# HPA006735,<br>RRID:AB_1852384 |
| α-TASOR (FAM208A) | Atlas Antibodies | Cat# HPA017142,<br>RRID:AB_1852382 |
| α-MPP8 | Proteintech | Cat# 16796-1-AP,<br>RRID:AB_2266644 |
| α- Beta Tububilin | Santa Cruz | Cat# sc-5274, RRID:AB_2288090 |
| α-P-smad2/3 | Cell Signaling Technology | Cat# 8828, RRID:AB_2631089 |
| α-Smad2/3 | Abcam | Cat# ab202445 |
| α-Anti-LINE-1 ORF1p | Abcam | Cat# ab216324,<br>RRID:AB_2921327 |
| α-Anti Flag M2 | Millipore sigma | Cat# <u>F3165</u>, RRID:AB_259529 |
| α-BrdU | BD Biosciences | Cat# 347580, RRID:AB_10015219 |
| α-CldU | Abcam | Cat# ab6326, RRID:AB_305426 |
| Donkey α-rabbit IgG (H+L)Antibody,<br>Alexa Fluor™ 647 | Invitrogen | Cat# A31573, RRID: AB_2536183 |
| Donkey α-mouse IgG (H+L)<br>Antibody, Alexa Fluor™488 | Invitrogen | Cat# A21202, RRID: AB_141607 |
| Donkey α-Goat IgG (H+L) Antibody,<br>Alexa Fluor™ 555 | Invitrogen | Cat# A-21432, RRID: AB_2535853 |

**Table 3.**
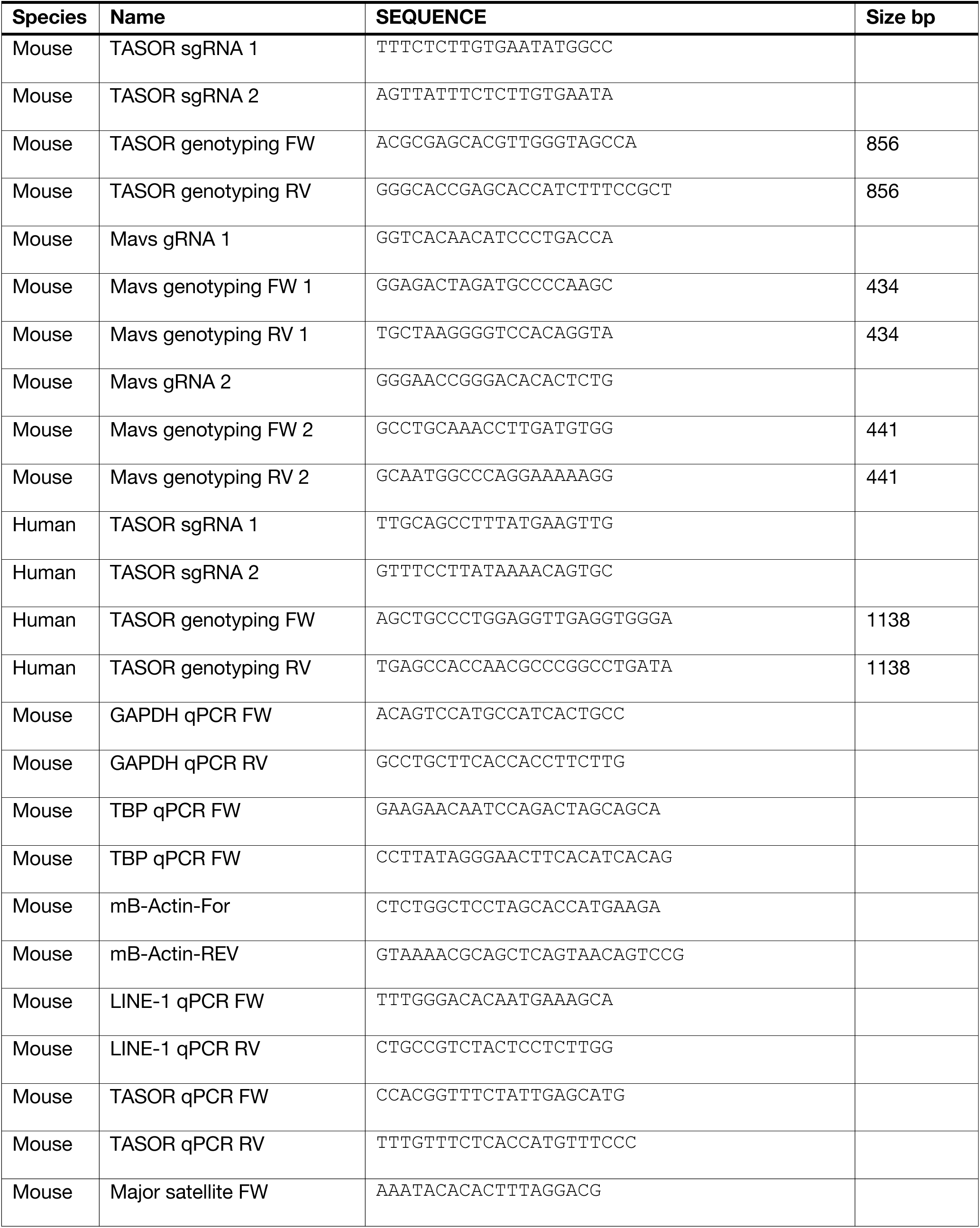
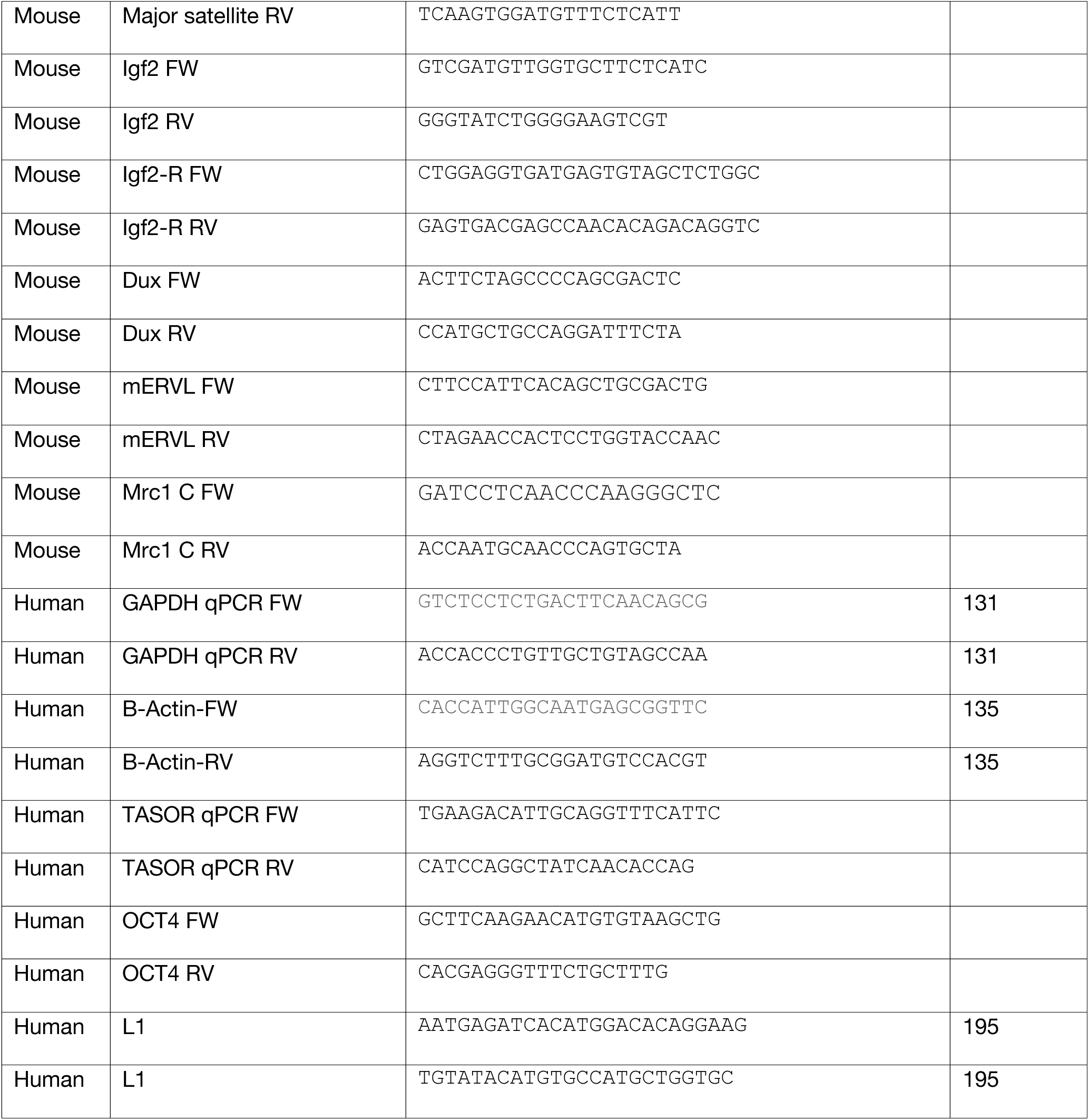
gRNAs and primers.

### Imaging

Phase contrast images were taken using a hybrid microscope (Echo Laboratories, CA) equipped with objective x2/0.06 numerical aperture (NA) air, x4/0.13 NA air, x10/0.7 NA air and 20x/0.05 NA air. Fluorescence imaging was performed on 8 or 96 well µ-siles (Ibidi) on a Nikon CSU-W1 spinning-disk super-resolution by optical pixel reassignment (SoRa) confocal microscope with objectives x4/0.13 NA, a working distance (WD) of 17.1nm, air; ×20/0.45 NA, WD 8.9–6.9 nm, air; ×40/0.6 NA, WD 3.6–2.85 nm, air.

### Imaging analysis

All Imaging experiments were repeated at least twice, with consistent results. In the figure captions, n denotes the number of biological repeats. Raw images were first processed in Fiji^110^ to create maximal intensity projection (MIP) and an export of representative images. Nuclear localized fluorescence intensity was computed for each cell in each field, and the value was then normalized to the DAPI intensity of the same cell. Intensity values of all cells were plotted as channel intensity over DAPI intensity for the same cell with mean ± s.d. GATA6 positive cells were selected and separated from negative cells using the spot colocalization tool. Epiblast cells were calculated as GATA6 negative spots. Trophectoderm and hypoblast cells as GATA6 positive spots.

### Western blotting

A minimum of 1×10^6 cells were harvested by centrifugation and lysed in RIPA lysis buffer (150mM NaCL, 1% Nonidet P-40, 0.5% Sodium deoxycholate (DOC), 0.1% SDS, 50mM Tris-HCL) supplemented with 1mM PMSF, 2mM MgCl_2_, 1x Halt complete protease inhibitor cocktail (Thermo Fisher Scientific) and 1x Halt phosphatase inhibitor cocktail (Thermo Fisher Scientific). Cell lysates were incubated with 10µl of benzonase (Sigma) per 100µl of RIPA buffer, for 15minutes at room temperature. Lysates were quantified using PIERCE BCA protein assay kit (Thermo Fisher Scientific) as per manufacturer instructions and absorbance was measured at 562nm using a SpectraMax iD3 plate reader (Molecular Devices). Protein concentrations were normalized to the lowest sample. Samples were denatured with Laemmli buffer (0.05M Tris-HCl at pH 6.8, 1% SDS, 10% glycerol, 0.1% β-mercaptoethanol) by boiling for 10 minutes. 30µg of total protein were resolved using SDS-PAGE followed by transfer to PVDF membranes. Transfer was visualized using Ponceau S staining solution (0.5 % w/v Ponceau S, 1% acetic acid). Membranes were cut and washed with TBS with 0.1% Tween (TBS-T) and blocking for 1 h with 5% MILK (BSA for phosphorylation specific antibodies) in TBS-T. Membranes were then incubated with the corresponding primary antibodies (Table 2). Immunoreactive bands were visualized using HRP conjugated secondary antibodies (Table 2), and incubated with chemiluminescence substrate (Pierce ECL western substrate, Thermo Fisher Scientific) and exposed to X-ray film or a ChemiDoc imaging system (BioRad).

### Teratoma tumor formation assay

Mouse ESCs were suspended in a 1:1 mixture of Matrigel and DMEM/F12 medium at a concentration of 1×10^7^ cells/mL. 100 µL of each cell mixture (1×10^6^ cells per tumor) was injected subcutaneously into the flanks of immunodeficient NOD/SCID mice. After 4 weeks, tumors were dissected, weighed, and fixed in 4% paraformaldehyde for 48 hours. Fixed tumor samples were submitted to the UT Southwestern Histopathology Core Facility for paraffin embedding and sectioning. Embedded and sectioned teratomas were stained with hematoxylin and eosin for tissue identification.

### Cell cycle analysis and Flow cytometry

Approximately 1×10^6 cells were treated with 10 µM EdU for 20 minutes at 37°C. Cells were harvested and fixed in ice-cold 96% methanol cells were permeabilized for 15 minutes using saponin and stained using the Click-iT Edu Alexa Fluor 488 flow cytometry assay kit (Thermo) as per manufacturing instructions, with DNA was counter staining with FxCycle Far Red with addition of RNAse (Thermo). Flow cytometry was performed using the appropriate unstained and single stain controls in a DBiosciences LSR II flow cytometer and analyzed using Flow Jo. Gating Strategy to determine cell cycle stages is shown in Figure S1G.

### IRF3 Dimerization assay

Cell pellets were resuspended in a modified RIPA buffer (50mM Tris Ph 8.0, 150mM NaCl, 1% NP40, 5% glycerol, 10mM sodium Fluoride, 0.4mM ETA, 1mM PMSF, 1xProtease inhibitor cocktail, 1xPhosphatase inhibitor cocktail), incubated on ICE for 30 min and centrifuge at 13,000xg for 10 min. Supernatant protein was quantified using the Pierce BCA quantification kit using a 1:2 dilution and absorbance at 563nm was detected in a spectrophotometer. Protein was flash frozen in liquid nitrogen and stored at −80°C. A 1.5mm non-denaturing 6% acrylamide/bis-acrylamide (29:1) gel without SDS at 4°C with the inner chamber buffer containing 25mM Tris-HCL pH 8.4, 192mM glycine and 1% deoxycholic acid in dH2O and the outer chamber containing 25mM Tris-HCl pH 8.4 and 192mM Glycine in dH2O was pre ran for 30 min on ice at constant 40 mA. 50µg of total protein was mixed with 2x loading dye (125mM Tris-HCL 6.8, 30% glycerol 0.1% Bromophenol blue in dH20) and ran at 40mA until the migration of the bromophenol blue exited the gel. The gel was washed for 15 minutes in 1x SDS-PAGE running buffer and transferred on ice to a PDVF membrane in 1x transfer buffer with 5% methanol at 375mA for 2 hours.

### 5-mC DOT blot assay

Dot blot analysis was made as described in Blaschke, et al^4^ with modifications, samples genomic DNA was purified from 1×10^6^ cells using the DNeasy blood & tissue Kit (Quiagen). DNA was eluted in 10mM Tris HCl pH 8 and quantified using a spectrophotometer. 2µg of DNA samples were denatured in 0.4M NaOH, 10mM EDTA at 100°C for 10 minutes, and neutralized by adding an equal volume of ice cold 2M ammonium acetate pH 7.0 and, and then serially diluted twofold. Nitrocellulose membranes were pre wetted in 6xSSC, diluted DNA samples were spotted on a nitrocellulose membrane using a Bio-Dot microfiltration apparatus (Bio-Rad). The blotted membrane was washed in 2× SSC buffer, dried at 80°C for 5min, and UV cross-linked at 120,000μJ/cm^2^. The membrane was then blocked in 5% milk in TBS-T for 1h at room temperature. Mouse anti-5-methylcytosine monoclonal antibody (Active Motif, 1:500) was added for 2h at room temperature. The membrane was washed for 10min three times in TBS-T, and then incubated with HRP-conjugated goat anti-mouse immunoglobulin-G (IgG) (Thermo, 1:10,000) for 1h at room temperature. The membrane was then washed for 10min three times in TBS-T and visualized with chemiluminescence substrate (Pierce ECL western substrate, Thermo Fisher Scientific) and a ChemiDoc imaging system (BioRad).

### Methylated DNA Immunoprecipitation and qPCR (meDIP qPCR)

Methylated DNA Immunoprecipitation was perfomed as described by Karpova, et al^111^, with modifications. Genomic DNA was purified from 1×10^6^ cells using the DNeasy blood & tissue Kit (Quiagen). Purified Genomic DNA was diluted to 100ng/µl in TE (10mM Tris, 1mM EDTA), 2.5µg in 62.5µl were sonicated for 6 cycles of 30 seconds on / 90 seconds off in 1.5ml tubes for 8 cycles at 4°C in a Bioruptor pico (sonication device (Diagenaode). Sonication size was confirmed via gel electrophoresis. DNA was denatured at 100°C for 10 minutes and quickly submerged in an ice water bath for 5 minutes. Immunoprecipitation was performed in 1x IP buffer (10mM sodium phosphate buffer pH 7.0, 1400mM NaCl, and 0.1% Triton X-100) with 1µg of antibody (Mouse anti 5mC, Active Motif, 39649 / Mouse IgG) per 1µg of DNA in a 100µl volume, at 4 °C overnight in a mixing platform. The following day 20µl of prewashed protein magnetic beads were added to each reaction, mixed and incubated at 4°C with overhead shaking. Beads were placed in a magnetic rack and washed x4 times with 200µl of 1x IP buffer and resuspended in 50µl of Proteinase K digestion buffer (50mM Tris-HCl pH 8.0, 10mM EDTA pH8.0 AND 0.1% SDS with 10µg/ml of proteinase K for 1h at 56°C with overhead shaking in a hybridization oven (UVP, Hybrilinker HL-2000). Beads were collected with a magnetic rack and supernatant was transferred to clean PCR tubes. DNA was the purified with AMPure XP beads (Beckam), briefly beads were mixed using a 1:1 ratio and incubated for 2 minutes and washed with 70% ethanol before eluting with 100µl of 10mM Tris-HCl pH 8.0. 8% of input was diluted 10 times and 2µl (4ng) were used for each qPCR reaction.

### Cloning of TASOR by overlapping PCR

TASOR was amplified by overlapping PCR, three primer sets were designed with overlapping sections,100ng of Genomic DNA was used per PCR reaction with primeSTAR GXL DNA polymerase (Takara Bio) using a touchdown PCR for the first 10 cycles from 72 to 60 followed by 35-40 cycles at the proper annealing temperature (Tm −2°C) and extension 68°C 30sec/Kb or 72°C 15sec/Kb and purified using a PCR purification KIT (Qiagen). Equimolar amounts of PCR products were mixed and a PCR was made with a primeSTAR GXL DNA polymerase (Takara Bio) without primers for the first 10 cycles using the following thermocycler conditions (95°C 3min, 98°C 10s, 60°C 30s, 68°C 5min, go to 2 15x, followed by the addition of the forward and reverse primers (0.5µM ea) and the reaction continued as a normal PCR for the next 20 cycles. Reaction products were gel purified and cloned into the expression vector via Gibson assembly. Vectors were sequenced using nanopore sequencing sanger sequencing at repetitive sites (Eurofins genomics).

### Auxin induced degradation of TASOR

Auxin-inducible degradation of TASOR was made using the auxin inducible degron 2 technology ^35^. For this mouse TASOR was cloned via Gibson assembly into a custom vector, expressing puromycin selection under a cytomegalovirus (CMV) enhancer fused to the chicken beta-actin promoter (CAG) promoter, a T2A sequence and TASOR with a C-terminal mini auxin-inducible degron (mAID) and a strep-strep-3xFlag tandem affinity purification (TAP). Oryza sativa TIR1 (OsTIR1) F74G was cloned downstream of the blasticidin resistance gene driven by a CAG promoter using a T2A sequence. Vectors were sequenced using nanopore sequencing Sanger sequencing at repetitive sites (Eurofins genomics). Cell lines were generated in a sequential manner via random integration and antibiotic selection with 5µg/ml of blasticidin and 1µg/ml of puromycin. Expression of TASOR and OsTIR1 was confirmed via qPCR and Flag immunofluorescence. Degradation was induced via the addition of 2µM 5ph-IAA, fresh media with 5ph-IAA was replaced every 12 hours. For mRNA half-live experiments, transcriptional inhibition was made with 5µg/ml of Actinomycin D

### Statistical analysis

All experiments were performed using two or more independent biological replicates. For cell cycle, q-PCR, imaging analysis and cell death assays, after verifying for the assumptions of equal variance and normality, P values were calculated using One-Way ANOVA with Tukey’s HSD. Unless otherwise indicated, error bars represent standard deviation. Analyses were performed with Prism (Graphpad).

### Visualization of RNA-seq and CUT&Tag data

The transcriptome and ChIP-seq datasets were visualized using Integrative Genomics Viewer (IGV, version 2.3.88 ^112^. Heatmaps and volcano plots were generated using R Statistical Software and the following R packages ggplot2, DESeq2m heatmap3, RcolorBrewer.

### CRISPR-Cas9 mediated gene knockout

CRISPR-Cas9 sgRNAs^25^ were design as previously descrived^113^, briefly all DNA sequences were manipulated using Benchling, sgRNAs were designed using Benchling with guide cleavage efficiency made with the WU-CRISPR tool^114^.Guides with minimal off targets and high cleavage efficiency were chosen. Each guide and its complementary sequence were ordered as synthetic 25nm oligos from (Thermo Fisher) with attached BbsI cloning sites: Sense: 5′–CACCGNNNNNNNNNNNNNNNNNNN–3′ and antisense: 3′–CNNNNNNNNNNNNNNNNNNNCAAA–5. Guides were cloned via golden gate assembly as described in Konnermann, S. et al^115^, with modifications. Briefly, 100pmol of each complementary oligo were phosphorylated using T4 PNK with T4 ligase buffer (contains ATP) for 30 minutes at 37°C, then oligos were annealed by denaturing at 95°C for 5 minutes and then slowly cooled down using a ramp of 5°C per minute up to 25°C. Phosphorylated and annealed oligos were diluted 1:10 and a Golden Gate reaction was setup with 1x rapid ligase (Roche), 10 units of BbsI enzyme, T7 DNA ligase (Roche) 25ng of backbone vector, reaction was run for 15 cycles of digestion at 37°C for 5 minutes and ligation at 20°C for 5 minutes. 2 µl of the golden gate reaction were transform into competent cells. After confirmation via sanger sequencing, maxipreps of sgRNAs were made with the Purelink Hipure plasmid maxiprep kit (Thermo) and DNA was eluted in 10mM Tris-HCl at a concentration of 2µg/µl. Cells were transfected using a NEPA21 electroporator (Nepa Gene) using 4 µg of total DNA per 1×10^6 cells. Conditions. Four “poring pulses” were applied (150 V, 3.0 ms, interval 50 ms, 10% voltage decay, polarity+), followed by 5 “transfer pulses” (5 V, 50 ms, interval 50 ms, 40% voltage decay; alternating + and − polarity). Cells were rapidly placed in pre warmed culture media to recover. Cells were FACS sorted 24 to 72h after transfection, the top 20% of selection marker positive cells was sorted and collected. Cells were plated at a density of 1000 cells per 9.6cm^2^. Single cell colonies were marked with an object maker (Nikon) and manually picked under a microscope in a laminar flow hood. 50 percent of recovered cells was lysed in 50µl of Quickextract DNA extraction solution (Biosearch) and genotyped via PCR. Positive clones were expanded and PCR again to ensure no integration of the CAS9 backbone.

### Mitotic Errors

Images were captured on a DeltaVision Ultra (Cytiva) microscope system equipped with a 4.2Mpx sCMOS detector. Fibers were acquired with a x100 objective (UPlanSApo, 1.4 NA) and 10×0.2μm z-section.

### DNA Fiber assay and analysis

To evaluate replication forks via DNA fibers^116^, exponentially growing cells were pulse-labeled for 20 minutes with 25 µM 5-iodo-2-deoxyuridine (I7125, Sigma-Aldrich), followed by a second 20-minute pulse with 250 µM 5-chloro-2-deoxyuridine (C6891, Sigma-Aldrich). The labeled cells were then washed twice with ice-cold 1X PBS, collected, and suspended at a concentration of 30,000 cells/ml. Subsequently, 30 µl of the suspension was centrifuged onto slides for 4 minutes at 800 rpm. After cytospinning, the slides were immersed in Lysis Buffer (0.5% SDS, 200mM Tris-HCl, 50mM EDTA) for 5 minutes, and DNA molecules were stretched using a homemade LEGO device. DNA fiber spreads were fixed in ice-cold Carnoy fixative for 10 minutes at room temperature and air-dried. Slides were rehydrated twice in water and incubated for 1 hour at room temperature in 2.5 M HCl. Afterward, the slides were rinsed twice in 1X PBS and blocked for 1 hour at room temperature in a blocking solution (1X PBS + 1% BSA + 0.5% Triton X-100 + 0.02% NaN3). The slides were then incubated in primary antibodies overnight at 4°C. The following primary antibodies were used at the indicated dilutions: 1:100 anti-BrdU (BDB347580, Becton Dickson) and 1:250 anti-CldU (ab6326, Abcam). The slides were then rinsed three times in 1X PBS and fixed in 4% paraformaldehyde in PBS for 10 minutes at room temperature. Afterward, they were rinsed twice in 1X PBS and incubated with 1:1,000 dilutions of Alexa Fluor-conjugated donkey anti-mouse or donkey anti-rat secondary antibodies (Invitrogen) for 2 hours at room temperature. Finally, the slides were washed twice in 1X PBS and mounted in ProLong Gold antifade mounting solution. Immunofluorescence images were captured on a DeltaVision Ultra (Cytiva) microscope system equipped with a 4.2Mpx sCMOS detector. Fibers were acquired with a x60 objective (PlanApo N 1.42 oil) and 1×0.2μm z-section. Quantitative image analyses were performed using Fiji (v.2.1.0/1.53c). locking Buffer.

### RNA Seq

RNA extraction was performed using a RNeasy Mini Kit (QIAGEN) using DNase treatment (QIAGEN). RNA was analyzed using a 2100 Bioanalyzer (Aglient Technologies). Libraries with unique adaptor barcodes were multiplexed and sequenced on an NovaSeq 6000 (paired-end, 150 base pair reads). Typical sequencing depth was at least 50 million reads per sample.

### RNA Seq analysis

Quality of datasets was assessed using the FastQC tool. Raw reads were adapter and quality trimmed using Trimgalore^117^. Reads were aligned to the mouse genome (mm10) with STAR^118^, using a custom GTF file which contained the NCBI RefSeq genes plus the addition of DFAM’s non-redundant repetitive element annotations^119^. Optical duplicate reads were filtered using Picard (http://broadinstitute.github.io/picard/). Samtools was used to filter out alignments with MAPQ < 30. Count matrices were generated using the featureCounts tool^120^. DESeq2 was used for the generation of normalized counts, log2FoldChange, and adjusted *p*-values^121^. baseMean was calculated as the mean of the normalized counts for samples present within a pairwise comparison. MA plots were generated using R and ggplot2^122^.

### CUT&Tag

CUT&Tag was performed as previously described^60^ with modifications. Briefly, 3×10^6 cells were harvested and resuspended in ice-cold nuclei extraction buffer (20mM HEPES KOH pH 7.9, 10mM KCl, 1% Triton X-100, 0.5 mM spermidine, EDTA free protease inhibitor cocktail (Roche)) at left on ice for 10 minutes. Nuclei were pelleted in a swinging bucket rotor at 1,300xg for 3 minutes, washed once with PBS and resuspended and cryopreserved in wash buffer 150 (20 mM HEPES pH 7.5, 150 mM NaCl, protease inhibitor cocktail (Roche), 0.5 mM Spermidine) with 10% DMSO, cryovials were placed in −80°C, using in CoolCell freezing containers (Corning) and then stored in liquid nitrogen until experiment. Nuclei were bound to CUTANA Concanavalin A Beads (Epicypher) for 15 min, then incubated with 50 μL Wash125 + 0.1% BSA, 2 mM EDTA, and 1 μL primary antibody targeting either H3K9me3 (Abcam, ab8898) or Flag (Cell Signalling, 2368T) overnight at 4°C. Nuclei were resuspended in 100 μL Wash125 + 1 μL secondary antibody at room temperature for 1 h. Nuclei were washed twice in 1 mL Wash125 (with no Spermidine), then resuspended in 200 μL Wash125 - Spermidine and + 0.2% formaldehyde for 2 min and then quenched with 50 μL 2.5 M glycine. Nuclei were washed once in 1 mL Wash350 (20 mM HEPES pH 7.5, 350 mM NaCl, 10 mM NaButyrate, 0.025% Digitonin, protease inhibitor cocktail (Roche), 0.5 mM Spermidine) then incubated in 47.5 μL Wash350 + 2.5 μL pAG-Tn5 (Epicypher 15-1017) for 1 h. Nuclei were washed twice in 1 mL Wash350, then resuspended in 300 μL Wash350 + 10 mM MgCl2 and incubated for 1 h at 37°C. Tn5 reaction was stopped with 10 μL 0.5 M EDTA, 3 μL 10% SDS, and 3 μL 18 mg/mL Proteinase K, briefly vortexed, then incubated at 55°C for 2 h to reverse crosslinks and release fragments. The fragments were then purified with phenol-chloroform and resuspended in 22 μL 1 mM Tris-HCl pH 8, 0.1 mM EDTA. The entire sample was amplified with Nextera i5 and i7 primers according to the Illumina protocol. The quality of the libraries was assessed using a D1000 ScreenTape on a 2200 TapeStation (Agilent) and quantified using a Qubit dsDNA HS Assay Kit (Thermo Fisher). Libraries with unique adaptor barcodes were multiplexed and sequenced on an NovaSeq 6000 (paired-end, 150 base pair reads). Typical sequencing depth was at least 50 million reads per sample.

### CUT&Tag analysis

Quality of datasets was assessed using the FastQC tool. Raw reads were adapter and quality trimmed using Trimgalore^117^. Trimmed reads were aligned to the mouse reference genome (mm10) with Bowtie2^123^ bowtie2 -q -R 3 -N 1 -L 20 -i S,1,0.50 --end-to-end --dovetail --no-mixed -X 2000). Multimapping reads were randomly assigned. Optical duplicate reads were filtered using Picard. Reads which mapped to the mitochondrial genome were removed with Samtools^124^ (samtools idxstats $sample.sorted.bam | cut -f 1 | grep -v chrM | xargs samtools view -b $sample.sorted.bam). Peak calling was performed with MACS2 software^125^ (--keep-dup all --nomodel -B -f BAMPE, --broad peakcalling was used for H3K9me3, whereas default narrow peaks were called for TASOR-FLAG and) and an FDR cutoff of 0.001 was applied to generate peak bedfiles. Peaks which intersected blacklisted high-signal genomic regions were removed. BigWig files were generated from merged bam files using deepTools^126^ and normalized to counts per million (CPM). Visualization of bigWigs was done in Integrative Genomics Viewer^112^. Intersections between different peak sets were made using BEDTools ^112^. Browser-style heatmaps and average profiles were generated using deepTools. Clustered heatmaps (ie Figure 3D) were generated using R and pheatmap^127^.

### ATAC-seq

The modified ATAC-sequencing protocol, Omni-ATAC was performed as previously described^128^. Briefly, 10^5^ cells were lysed with resuspension buffer (Tris 10 mM, pH 7.4, 10 mM NaCl, 3 mM MgCl2, 0.1% NP-40, 0.1% Tween-20, and 0.01% Digitonin) and nuclei were collected for tagmentation at 37 °C for 30 minutes (Illumina Tagment DNA Enzyme and Buffer Small Kit). The reaction was immediately purified using Qiaquick PCR Purification Kit (Qiagen) and eluted in 20 μl water. Eluted DNA was amplified using NEBNext Ultra II PCR Master Mix (NEB) and purified using AMPure XP beads. Libraries with unique adaptor barcodes were multiplexed and sequenced on an NovaSeq 6000 (paired-end, 150 base pair reads). Typical sequencing depth was at least 50 million reads per sample.

### ATAC-Seq Analysis

ATAC-seq data was processed as described above for CUT&Tag.

### TOBIAS analysis

For TOBIAS analysis, replicate bam files were merged using Samtools. TOBIAS ATACorrect and ScoreBigWig were used to generate scored bigWig files for each merged sample. BINDetect was then used to generate pairwise differential binding scores between samples for each expressed JASPAR motif. For analysis of differential binding scores specifically in promoters, BINDetect was restricted using option --output-peaks to regions of interest, ie specific repetitive element subfamilies using bedfiles generated from Dfam’s dfam’s non-redundant hits files (mm10.nrph.hits.gz) or the non-repetitive genome.

## Author Contributions

C.A.P-A, R,O., L.B., and J.W. conceptualized, designed, analyzed, and interpreted the experimental results. A.M performed mitotic error and DNA fiber analysis. Y.H generated the MAVS TASOR double knockout lines. E.B, Y.W, M.S, performed teratoma assays and mouse work. A.P performed experimental procedures and data collection. R.O. performed epigenetic profiling and bioinformatic data processing. C.A.P-A and D.S performed molecular cloning. P.L., L.B. and J.W. supervised the study. C.A.P-A., A.P, R.O., L.B. and J.W. wrote the manuscript with inputs from all authors.

## Acknowledgements

We would like to thank the members of the Wu, Banaszynski, Moazed and Tagliabracci for discussions throughout this project. We would also like to help inputs from Harleen Saini, Vincent Tagliabracci, Joshua Mendell, Zhijian ‘James’ Chen and Eric Olson. We want to acknowledge the help from Krzysztof Pawlowski in the bioinformatic discovery. We would like to acknowledge the help from Dr Angela Mobley (UT Southwestern Flow cytometry core), Dr. Marcel Mettlen (UT Southwestern Quantitative Light Microscopy Core Facility) Dr. Andrew Lemoff (UT Southwestern Proteomics Core Facility), and Drs. Chad Brautigam and Dr. Shih-Chia (Scott) Tso (UT Southwestern Molecular Biophysics Core). L.B. is a Virginia Murchison Linthicum Scholar in Medical Research and funded by NIH (GM124958 and HD109239), the American Cancer Society (134230-RSG-20-043-01-DMC), and the Welch Foundation (I-2025). J.W. is a New York Stem Cell Foundation–Robertson Investigator and Virginia Murchison Linthicum Scholar in Medical Research and funded by CPRIT (RR170076), NIH (GM138565-01A1 and OD028763), Welch (854671). The Nikon SoRa spinning disk microscope was purchased by the UTSW quantitative light microscopy core with a Shared Instrumentation grant from NIH award: 1S10OD028630-01 to Katherine Luby-Phelps.

